# Structure-Based Protein Function Prediction using Graph Convolutional Networks

**DOI:** 10.1101/786236

**Authors:** Vladimir Gligorijevic, P. Douglas Renfrew, Tomasz Kosciolek, Julia Koehler Leman, Daniel Berenberg, Tommi Vatanen, Chris Chandler, Bryn C. Taylor, Ian M. Fisk, Hera Vlamakis, Ramnik J. Xavier, Rob Knight, Kyunghyun Cho, Richard Bonneau

## Abstract

The large number of available sequences and the diversity of protein functions challenge current experimental and computational approaches to determining and predicting protein function. We present a deep learning Graph Convolutional Network (GCN) for predicting protein functions and concurrently identifying functionally important residues. This model is initially trained using experimentally determined structures from the Protein Data Bank (PDB) but has significant de-noising capability, with only a minor drop in performance observed when structure predictions are used. We take advantage of this denoising property to train the model on > 200,000 protein structures, including many homology-predicted structures, greatly expanding the reach and applications of the method. Our model learns general structure-function relationships by robustly predicting functions of proteins with ≤ 40% sequence identity to the training set. We show that our GCN architecture predicts functions more accurately than Convolutional Neural Networks trained on sequence data alone and previous competing methods. Using class activation mapping, we automatically identify structural regions at the residue-level that lead to each function prediction for every confidently predicted protein, advancing site-specific function prediction. We use our method to annotate PDB and SWISS-MODEL proteins, making several new confident function predictions spanning both fold and function classifications.

---

Proteins fold into 3-dimensional structures to carry out a wide variety of functions within the cell^1^. Even though many functional regions of proteins are disordered (lack a well defined ensemble average structure), the majority of protein domains in natural proteins fold into specific and ordered three-dimensional conformations ^2–6^. In turn, the structural features of proteins determine a wide range of functions: from binding specificity, forming structures like cytoskeletal elements within the cell, to catalysis of biochemical reactions, transport, and signal transduction. There are several widely used classification schemes that organize these myriad protein functions including: the Gene Ontology (GO) Consortium^7^, Enzyme Commission (EC) numbers^8^, Kyoto Encyclopedia of Genes and Genomes (KEGG)^9^, and others. GO, for example, classifies proteins into hierarchically related functional classes organized into 3 different ontologies: Molecular Function (MF), Biological Process (BP) and Cellular Component (CC), to describe different aspects of protein functions.

The advent of efficient low-cost sequencing technologies and advances in computational methods (e.g. gene-finding) have resulted in a massive growth in the number of sequences available in key protein sequence databases like the UniProt Knowledgebase (UniProtKB)^10^. UniProt currently contains over 100 million sequences, only ∼0.5 million (0.5%) of which are manually annotated (UniProtKB/Swiss-Prot). Due to considerations of scale, experimental design, and the cost of experimentally verifying a function, it is safe to posit that most proteins with unknown function (i.e., hypothetical proteins) are unlikely to be experimentally characterized. Understanding the functional roles and studying the mechanisms of newly discovered proteins is one of the most important biological problems in the post-genomic era. In parallel to the growth of sequence data, the advent of experimental as well as computational techniques in structure biology has made the three-dimensional structures of many proteins available^11–18^. The Protein Data Bank (PDB)^19^, a repository of three-dimensional structures of proteins, nucleic acids, and complex assemblies, has experienced significant recent growth, reaching over 150,000 entries. Large databases of comparative models (e.g., SWISS-MODELs where structure is mapped to query sequences via alignment) also provide valuable resources for studying structure-function relationships ^13, 20^.

To address the sequence-function gap many computational methods have been developed that aim to predict protein function for whole protein genes. A related body of work is also directed at the related problem of predicting function in a site- or domain-specific manner (methods that automatically generate functional hypothesis linked to residues, regions or domains)^21–24^. Traditional machine learning classifiers, such as support vector machines, random forests, and high-dimensional statistical methods like logistic regression have been used extensively for the protein function prediction problem, and have established that integrative prediction schemes can outperform homology-based function transfer^25, 26^ and that integration of multiple gene- and protein-network features typically outperform sequence based features (although network features are often incomplete or unavailable). Systematic benchmarking efforts, such as the Critical Assessment of Functional Annotation (CAFA1^27^, CAFA2^28^ & CAFA3^29^) and MouseFunc^30^, have also played a key role in the development of these methods and have shown that integrative machine learning and statistical methods outperform traditional sequence alignment-based methods (e.g., BLAST)^26^. However, the performance of these methods is typically strongly affected by the quality of manually-engineered features constructed from either sequence, biological networks or protein structure (features that rely heavily on heuristics that in turn require domain-expert knowledge, and in some cases unstable assumptions, thresholds and preprocessing pipelines)^31^. Here, we focus on methods that can take as inputs sequence and features that are readily derived from sequence (such as predicted structure) and do not focus on, or compare to, the many methods that rely on protein networks like *GeneMANIA*^32^, *Mashup*^33^, *DeepNF*^34^, and other integrative network prediction methods. We focus our study in this way to present a method that can be applied to very large volumes of sequences where many proteins are from unknown organisms and thus lacking the required network data. In this way we aim to address the critical need for function and structure annotation methods in larger-scale metagenomic contexts.

In the last decade, deep learning approaches have achieved unprecedented improvements in performance across a broad spectrum of problems ranging from learning protein sequence embeddings for contact map prediction^35^ to predicting protein structure^36, 37^ and function^38^. In particular, Convolutional Neural Networks (CNN)^39^, the state-of-the-art in computer vision, have also shown tremendous success in addressing problems in computational biology. These machine learning methods have enabled task-specific feature extraction directly from protein sequence (or the corresponding 3D structure) overcoming the limitations of feature-based Machine Learning (ML) methods. The majority of sequence-based protein function prediction methods use 1D CNNs, or variations thereof, that search for recurring spatial patterns within a given sequence and converts them hierarchically into complex features using multiple convolutional layers. Recent work has employed 3D CNNs to make predictions and extract features from protein structural data^40, 41^. Although these works demonstrate the utility of structure features, storing explicit 3D representations of protein structure at high resolution is not memory efficient, since most of the 3D space is unoccupied by protein structure. We experiment here with more efficient representations for the protein structure. Geometric deep learning methods^42, 43^, and more specifically Graph Convolutional Networks (GCNs)^44^, have offered a way to overcome these limitations by generalizing convolutional operations on more natural graph-like molecular representations. GCNs have shown tremendous success in various problems ranging from learning useful molecular fingerprints^45^, to predicting biochemical activity of drugs^46^, to classifying protein interface prediction^47^.

Here, we describe a method, *DeepFRI*, based on GCNs for functionally annotating protein sequences and structures that outperforms current methods and scales to the size of current repositories of sequence information (see Fig. 1). Our model is a two stage architecture that takes as input a protein sequence and structure (represented as graphs derived from amino acid interactions in the 3D structure) and outputs probabilities for each function. The first stage of the method is a self-supervised a Long Short-Term memory Language Model (LSTM-LM)^48^ pretrained on a set of protein domain sequences from the protein families database (Pfam)^49^ (see Methods), and is used for extracting residue-level features from PDB sequences. The second stage is a graph convolutional network that uses a deep architecture to propagate these residue-level features between residues that are proximal in the structure and construct final protein-level feature representations that we show are useful for protein function prediction below.

**Figure 1:**
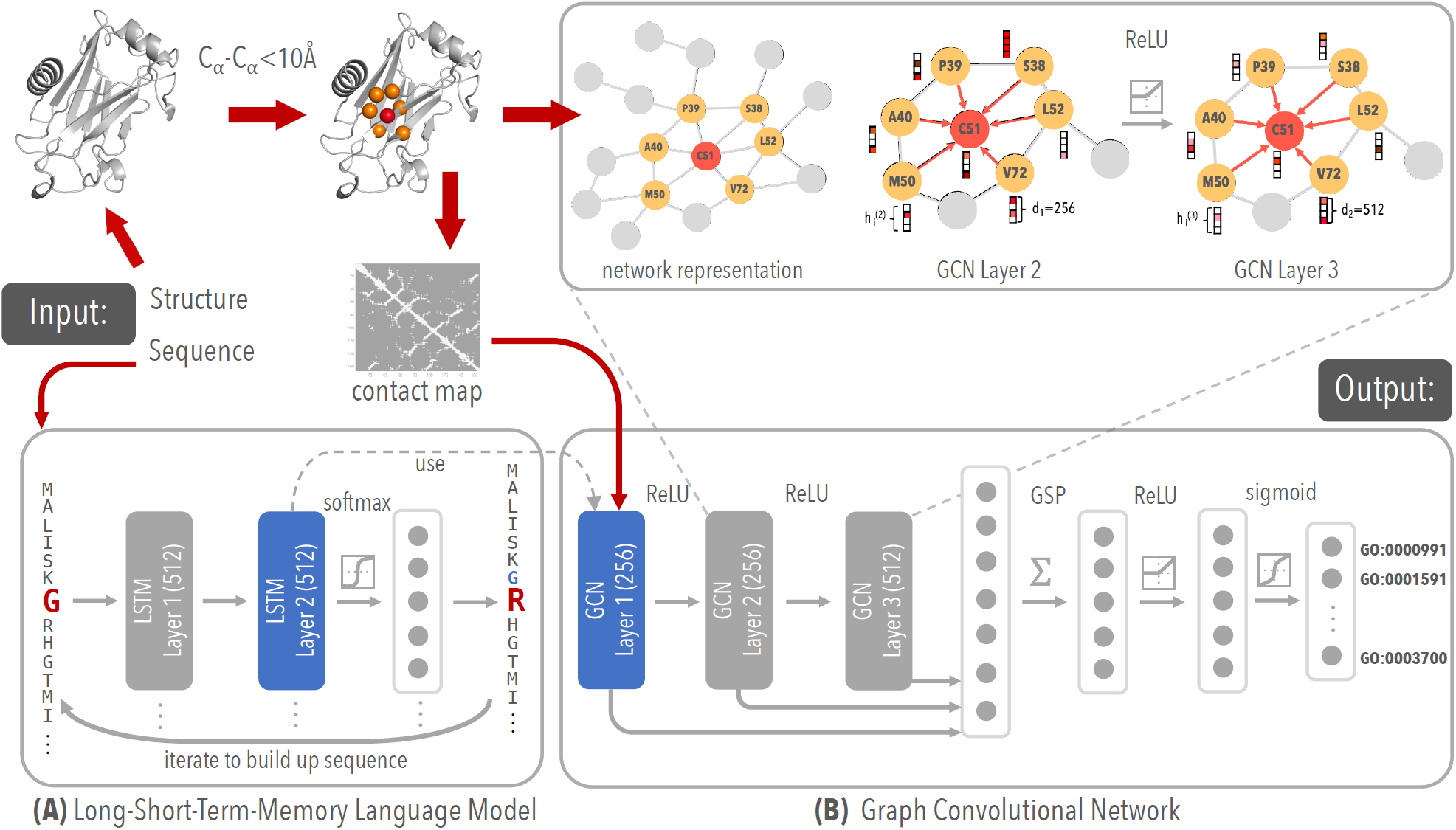
Schematic method overview. (A) LSTM language model, pretrained on ∼10 million Pfam protein sequences, used for extracting residue level features of PDB sequence. (B) our GCN with 3 graph convolutional layers for learning complex structure–to–function relationships.

For learning sequence features we use a LSTM-LM pretrained on a corpus of around 10 million protein domain sequences from Pfam^49^ (see Supplemental Methods). Our LSTM-LM is trained to predict an amino-acid residue in the context of its position in a protein sequence. Using features from a pre-trained, task-agnostic, LSTM-LM as input to classification tasks has demonstrated tremendous success in many Natural Language Processing (NLP)^50^ and biological problems^35^. During the training of the GCN the parameters of the LSTM-LM are frozen; i.e., the LSTM-LM stage is only used as a sequence feature extractor. The residue-level features constructed for sequences, together with contact maps, are used as an input for the second stage of our method. We show that such features can significantly boost performance of the GCN in function prediction task (see Supplementary Fig. 2A). Using LSTM-LM features together with contact maps we show that our method outperforms sequences-only state-of-the-art methods.

Each layer of the graph convolution stage takes both an adjacency matrix and the residue-level embeddings described above, and outputs the residue-level embeddings in the next layer. The GCN protein representation is obtained by concatenating features from all layers of this GCN into a single feature matrix and subsequently fed into two fully connected layers to produce the final protein function predictions for all terms (see Methods). This choice of architecture leads to the main advantage of our method, that it convolves features over residues that are distant in the primary sequence, but close to each other in the 3D space, without having to learn these functionally relevant proximities from the data. Such an operation, implemented here using graph convolution, leads to better protein feature representations and ultimately to more accurate function predictions. The effect of long-range connections on predictive performance of our method is shown below and in Supplementary Fig. 2B.

## Evaluating our method on experimental and predicted structures

We train different models to predict GO terms (one model for each branch of the GO) and EC numbers. First, we systematically investigate the performance of our method trained only on experimentally determined, high-quality, structures from the PDB. Then, we examine the performance when homology-based predicted structures from SWISS-MODEL are included in the training procedure. GO term and EC number annotations for PDB and SWISS-MODEL structures are retrieved from SIFTS ^51^ and UniProtKB/Swiss-Prot repositories, respectively (see Methods for data collection and preprocessing). We carefully explore the utility of using predicted structure (by multiple methods) for both training and prediction of newly observed sequences and find that the use of SWISS-MODEL structures during the training greatly improves model comprehension and accuracy.

We first explore how our method, trained on experimental PDB structures, tolerates errors in predicted structures by examining its performance on predicted structures obtained from SWISS-MODEL^13^ and other *de novo* protocols (see Fig. 2A). We use both Rosetta macromolecular modeling suite^52^ and protein contact predictions from DeepMetaPSICOV contact predictor (DMPfold)^12^ to fold sequences of ∼800 experimentally annotated PDB chains and obtain the lowest energy decoy from folding. We construct two kinds of *C*_*α*_ − *C*_*α*_ contact maps for each PDB chain – one from its experimental (i.e., *NATIVE*) structure and one from the lowest-energy (i.e., *LE*) decoy (see Methods). Our model exhibits higher performance than that of the CNN-based method (*DeepGOPlus*) even when accounting for errors in predicted contact maps from the full range of widely adopted methods tested (Fig. 2A). Even though Rosetta- and DMPfold-predicted structures often result in noisy contact maps, the fact that the performance of our method on the predicted LE structures is not drastically impaired (Fig. 2A, left) can be attributed to the high denoising ability of the GCN implied by high correlation between GCN features extracted from NATIVE and LE contact maps (see Supplementary Fig. 3).

**Figure 2:**
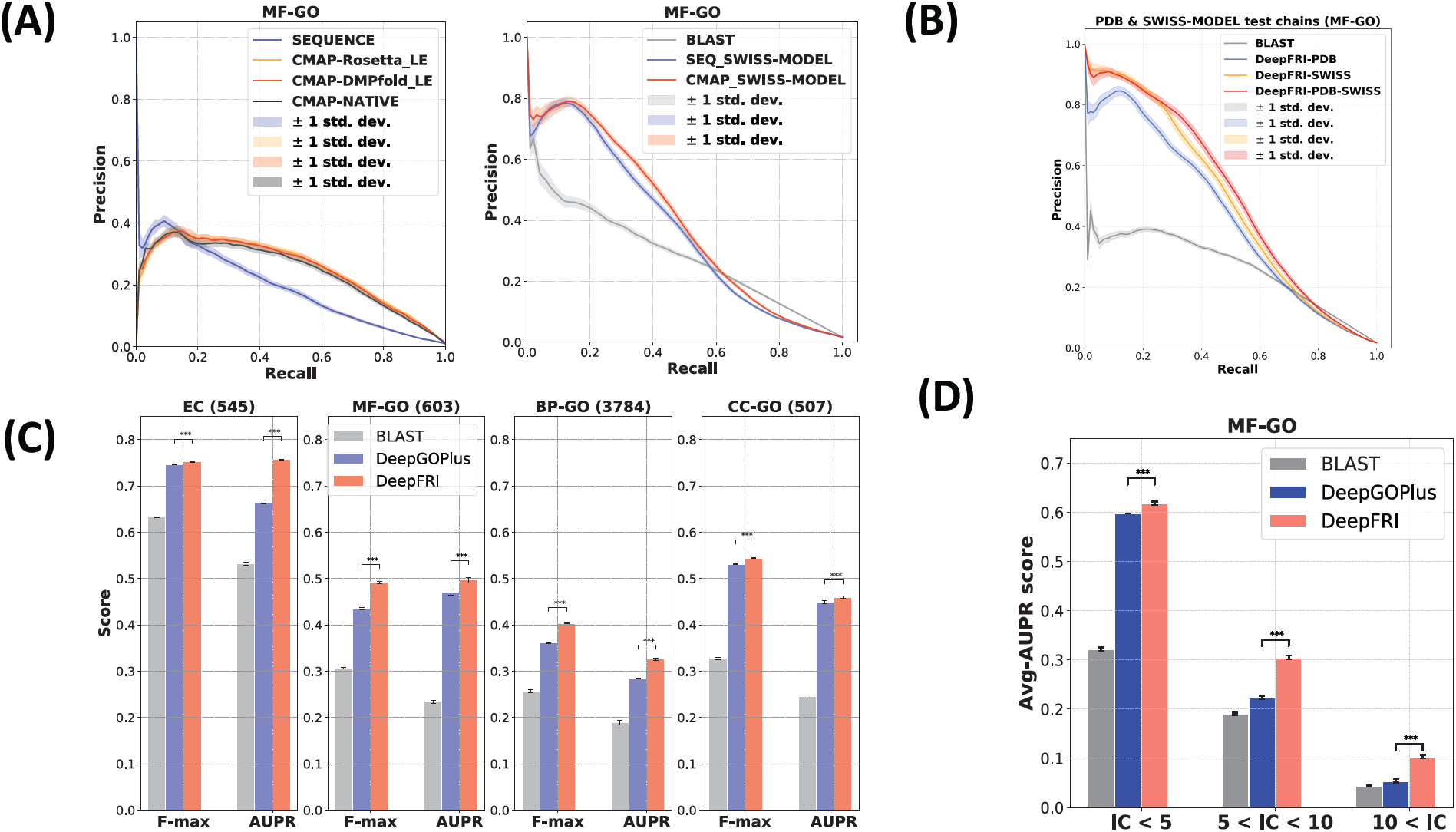
Improved performance on experimental and predicted structures. (A) (left) Precision-recall curves showing performance on ∼800 annotated PDB chains obtained using the CNN-based method (*DeepGOPlus*) applied to sequences (blue), our GCN-based method (*Deep-FRI*) applied to NATIVE protein structures (black), Rosetta-predicted lowest energy (LE) structures (yellow) and DMPfold lowest energy (LE) structures (red). All models are trained only on experimental PDB structures. (right) Precision-recall curves representing the performance of our method, trained only on PDB experimental structures, and evaluated on SWISS-MODEL homology-predicted structures (red); performance of *DeepGOPlus* trained on PDB sequences and BLAST baseline are shown in blue and grey, respectively. The curves are averaged over prediction results from 100 bootstraps of the test set; (B) Average performance of our *DeepFRI* method trained on PDB (blue), SWISS-MODEL (yellow) and both PDB and SWISS-MODEL (red) structures in comparison to *BLAST* baseline (grey). (C) F-max and AUPR scores, summarized over all proteins and GO terms, respectively, computed on the test set comprised of PDB and SWISS-MODEL chains chosen to have ≤ 40 % sequence identity to the sequences in the training set. Numbers in brackets indicate the number of GO terms in different ontologies. (D) Average AUPR performance of *DeepFRI* on MF-GO terms of different specificities in comparison to CNN-based architecture (*DeepGOPlus*) and *BLAST* baseline. The results are averaged over 100 bootstraps of the test set. Asterisks indicate where the performance of *DeepFRI* is significantly better than *DeepGOPlus* (rank-sum *p − val <* 0.001).

Second, we examine the possibility of including the predicted structures into the model training procedure. Having a large number of diverse protein structures in the training set is an important prerequisite for more accurate and robust performance of our deep learning-based method. To this end, we combine ∼40k non-redundant experimental structures from the PDB and ∼200k non-redundant homology-predicted structures from the SWISS-MODEL repository. Inclusion of SWISS-MODEL predicted structures not only results in more training examples, and consequently in more accurate performance (with AUPR of 0.501 ± 0.007 in comparison to AUPRs of 0.436 ± 0.007, 0.481 ± 0.006 of the models trained only on PDB and only on SWISS-MODEL chains, respectively; see Fig. 2B), but it also results in a larger GO term coverage, especially in the number of very specific, rarely-occurring GO terms (Information Content, IC > 10; Supplementary Fig. 4). Comparing the performance of our model with the CNN-based method, *DeepGOPlus*^38^, that operates only on sequences, and BLAST baseline (AUPR of 0.269 ± 0.004, Fig. 2B) we observe that our method benefits greatly from predicted structures.

For comparison to prior studies, we evaluate the performance of our model in comparison to other methods using commonly accepted measures also employed in the CAFA1 challenge ^27^: protein-centric maximum F-score (F-max) and term-centric Area Under Precision-Recall (AUPR) curve (see Supplementary Material). The AUPR and F-max scores are averaged over all GO terms and all proteins in the test set, respectively. The performance of our method for each branch of GO is shown in red in Figure 2C. The performance is compared to 1) *BLAST* baseline (grey) used in the CAFA1^27^, CAFA2^28^ & CAFA3^29^ function blind challenges, and to 3) *DeepGOPlus*, a state-of-the-art CNN-based method (blue) (see Methods for the architecture details). Comparison to standard feature engineering-based, SVM-based method, *FFPred*, is shown in Supplementary Fig. 5. Given that *FFPred* is limited in the number of GO terms for which it can make predictions (and also it cannot predict EC numbers), we only show the result averaged over a subset of GO terms common to all methods. It is important to note that different methods encompass different subsets of the GO-term vocabulary and that a key advantage of using comparative models in training is the increase in the size of the vocabulary encompassed by our method; for this reason the number of GO terms is indicated throughout each figure. Our method substantially outperforms both CNN-based methods and BLAST on the EC numbers (only the most specific EC numbers are considered in the training, i.e., leaf nodes in the EC tree), and GO terms in all three branches.

We explored the performance of our method on individual GO terms. We observe that for the majority of MF-GO terms, our method outperforms the sequence-only CNN method, indicating the importance of structure features in improving performance (see also Supplementary Fig. 6). Our method performs better than sequence-only CNN and the CAFA-like BLAST baseline for more specific MF-GO with fewer training examples (see Fig. 2D). Our method outperforms CNN on almost all GO terms with average PDB chain length ≥ 400 (see Supplementary Fig. 6), illustrating the importance of encoding distant amino-acid contacts via the structure graph. This demonstrates the superiority of graph convolutions over sequence convolutions in constructing more accurate protein features when key functional sites are composed of distal sequence elements (as is the case for higher contact order proteins)^53^. Specifically, in the case of long protein sequences, a CNN with reasonable filter lengths, would most likely fail to convolve over residues at different ends of the long sequence, even after applying multiple consecutive CNN layers; whereas, graph convolutions applied on contact maps would, in 3 layers or less, access feature information from the complete structure.

## From protein-level to region-level predictions

Many proteins carry out their functions through spatially clustered sets of important residues (e.g., active sites on an enzyme, ligand-binding sites on a protein, protein-protein interactions); this is especially the case for site-specific functions in the Molecular Function branch of the GO, and less true for terms encoded in GO Biological Process terms. Designing ML methods for identifying such functional residues have been the subject of many recent studies^21–24^. Much recent work in ML has provided several new approaches for localizing signal to regions of the input feature space that lead to a given positive prediction, providing a means of interpreting decisions made by neural networks^54, 55^. In computer vision these methods determine the regions of images that lead to positive object classifications; in NLP these methods lead to identification of sub-regions of documents^56^. Recent work in computer vision uses gradient-weighted Class Activation Maps (grad-CAMs) on trained CNN-based architectures^57^ to localize the most important regions in images relevant for making correct classification decisions^57^. We propose an approach adapted for GCNs for detecting functional regions in proteins, Deep Functional Residue Identification (*Deep-FRI*) that provides the ability to interpret predictions by analysing the relative importance of features during the training and during prediction. For each protein, our *DeepFRI* grad-CAM detects GO term-specific structural sites by identifying residues relevant for making accurate GO term prediction. We use grad-CAMs^57^, adapted for post-training analysis of GCNs, to determine a region of a protein that leads to the correct prediction of its GO function. For each GO term, the grad-CAM technique generates an activation map over the input data, in our case a sequence of residues and the contact or distance map, indicating the importance of each residue to the GO term classification decision (see an example of grad-CAM and its corresponding heatmap over sequence in Fig. 3A, right). It does so by first computing the contribution of each graph convolutional feature map of the model (trained on the MF-GO dataset) to the GO term prediction, and then by summing the feature maps with positive contributions to obtain a final residue-level activation map (see Methods).

**Figure 3:**
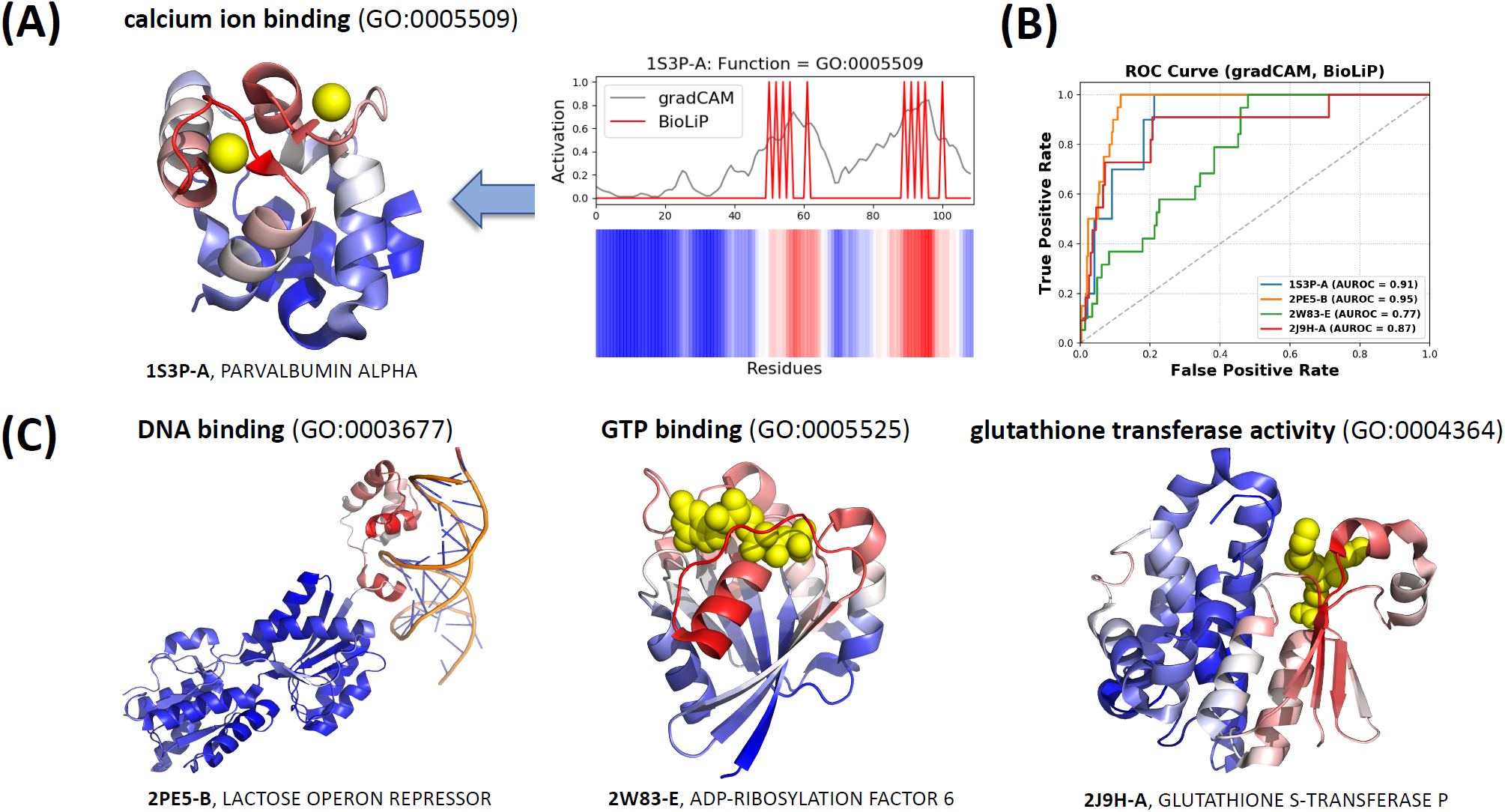
Automatic mapping of function prediction to sites on protein structures. (A) An example of the gradient-weighted class activation map for ‘Ca Ion Binding’ (right) mapped onto the 3D structure of rat alpha-parvalbumin (PDB Id: 1S3P), chain A (left), annotated with *calcium ion binding*. The two highest peaks in grad-CAM activation profile correspond to calcium binding regions. (B) ROC curves showing the overlap between grad-CAM activation profiles and binding sites, retrieved from the *BioLiP* database, computed for the PDB chains shown in panel C. (C) Examples of other PDB chains annotated with *DNA binding, GTP binding* and *glutathione transferase activity*. All residue are colored using gradient color scheme to match the grad-CAM activity profile, with more salient residues highlighted in red and less salient residues highlighted in blue. No information about co-factors, active sites, or site-specificity was available to the model during training.

For site-specific functions, this method identifies correct function regions and we provide several examples where we automatically and correctly identify functional sites for several functions where binding sites are known (see Fig. 3). Fig. 3A shows the grad-CAM identified residues for a *calcium ion binding (GO:0005509)* of alpha-parvalbumin protein (PDB id: 1S3P). The two highest peaks in the grad-CAM profile correspond to the binding regions in the 3D structure of the protein (Fig. 3A, left). Indices of the calcium binding residues of the 1S3P protein were retrieved from the BioLiP database^58^ and compared to the residues identified by our method. ROC curve computed between the binary profile representing binding sites from BioLiP (shown in red) and the grad-CAM profile (shown in grey) in Fig. 3A, right are depicted in Fig. 3B for several proteins. High area under the ROC curve indicates high correspondence between annotated binding sites and our predictions. Similar correspondence with BioLiP is observed for several other functions including *DNA binding (GO:0003677), GTP binding (GO:0005525)* and *glutathione transferase activity (GO:0004364)* (see Fig. 3C and their corresponding ROC curves in Fig. 3B). Our study of grad-CAMs against BioLiP database reveals that the highest performing group of GO terms are related to functions with known site-specific mechanisms or site specific underpinnings. We depict examples (with high AUROC scores) for which grad-CAMs correctly identify binding regions. We show that, for various GO terms, these functional sites correspond to known binding sites, conserved functional regions or experimentally characterized active sites (see Supplementary Fig. 7 and Supplementary Figs. 8-16). Interestingly, our model is not explicitly trained to predict functional sites, but instead such predictions stem solely from the grad-CAM analysis of the graph convolution parameters of the trained model; thus, the ability of the method to correctly map functional sites supports our argument that the method is general and capable of predicting functions in a manner that transcends sequence alignment. Performing such analysis for identifying functional sites is also very efficient as it does not require any further training or modification of the model’s architecture. The site-specificity afforded by our function predictions is very valuable, especially in the case when predicting functions of poorly studied, unannotated proteins. Site-specific predictions provide first insights into the correctness of predictions and frames follow-up validation experiments: for example, using genetics or mutagenesis to test site specific predictions.

We also evaluate the performance of our method in a more “realistic” scenario by using a *temporal holdout* validation strategy similar to the one in CAFA ^27–29^. That is, we composed a test set of PDB chains by looking at the difference in GO annotations of the PDB chains in the SIFTS ^51^ database between two releases separated by ∼6 months time period – realises 2019/06/18 and 2020/01/04. We identify ∼3,000 PDB chains that did not have any annotations in the 2019 SIFTS release and gain new annotations in the 2020 SIFTS release (see Methods). By using the annotations from the 2020 SIFTS release, we evaluated the performance of our method on these newly annotated PDB chains. We show that our method significantly outperforms both *BLAST* and *DeepGOPlus* (see Supplementary Fig. 17). Furthermore, we highlight particular examples of PDB chains with GO terms correctly predicted by our method for which both *BLAST* and *DeepGOPlus* are failing to produce any meaningful predictions, indicating again the importance of structure information (see Supplementary Fig. 17).

## Unannoated PDB and SWISS-MODEL chains

There are a large number of high quality protein structures in both the PDB and SWISS-MODEL that lack functional annotations, or that have only high-level or incomplete annotation in the databases we used for training and testing our models. For example, our analysis of the June 2019 release of the SIFTS database^51^ reveal that around 20,000 non-redundant, high quality PDB chains currently lack GO term annotations. Similarly, around 13,000 SWISS-MODEL chains also lack Swiss-Prot GO term annotations. Interestingly, even though these PDB chains lack actual GO term annotations, we found that many have additional site-specific functional information information present/evident in their PDB files (e.g. presence of ligands, co-factors, metals, DNA and RNA). We focus on these cases, as their function predictions can be verified and discussed in depth (our larger set of predictions, including many for truly unknown PDB chains, is provided in the Supplementary Files 1). For example, there are a number of metal ion-bound PDB chains, with known binding residues in BioLip^58^, but with missing *metal ion binding (GO:0046872)* GO term annotations. In other cases, the function, albeit missing in SIFTS, is directly implied in the name of the protein (e.g., a zinc finger protein without *zinc ion binding (GO:0008270)* annotation). Here, we apply our method to these “unannotated” PDB chains, as a part of blind experiment, to evaluate the chain-level predictions of our method and also the residue-level predictions of the grad-CAM approach. We also apply our method on SWISS-MODEL chains.

We present Supplementary Files 1 & 2 with all high confidence predictions for the PDB and SWISS-MODEL chains produced by our method. In Fig. 4 (panels A & B), we show their statistics, with the total number of PDB and SWISS-MODEL chains predicted to be annotated with all and more specific (IC > 5) GO terms. Some interesting “unannotated” PDB chains with known ligand-binding information include *4 iron, 4 sulfur cluster binding (GO:0051539)* of a Fe-S-cluster-containing hydrogenase protein (PDB id: 6F0K), is shown in Fig. 4C. Iron-sulfur clusters are shown to be important in oxidation-reduction reactions of electron transport and our method can accurately predict their binding sites as shown by the corresponding ROC curve computed between the predicted grad-CAM profile and known 4Fe4S cluster binary binding profile retrieved from BioLiP. Another example includes predicted *DNA binding (GO:0003677)* and *metal ion binding (GO:0046872)* of the zinc finger protein (PDB Id: 1MEY) with predicted grad-CAM activity mapped onto the same 3D structure and validated by experimental density for both DNA and metal as shown in Fig. 4D.

**Figure 4:**
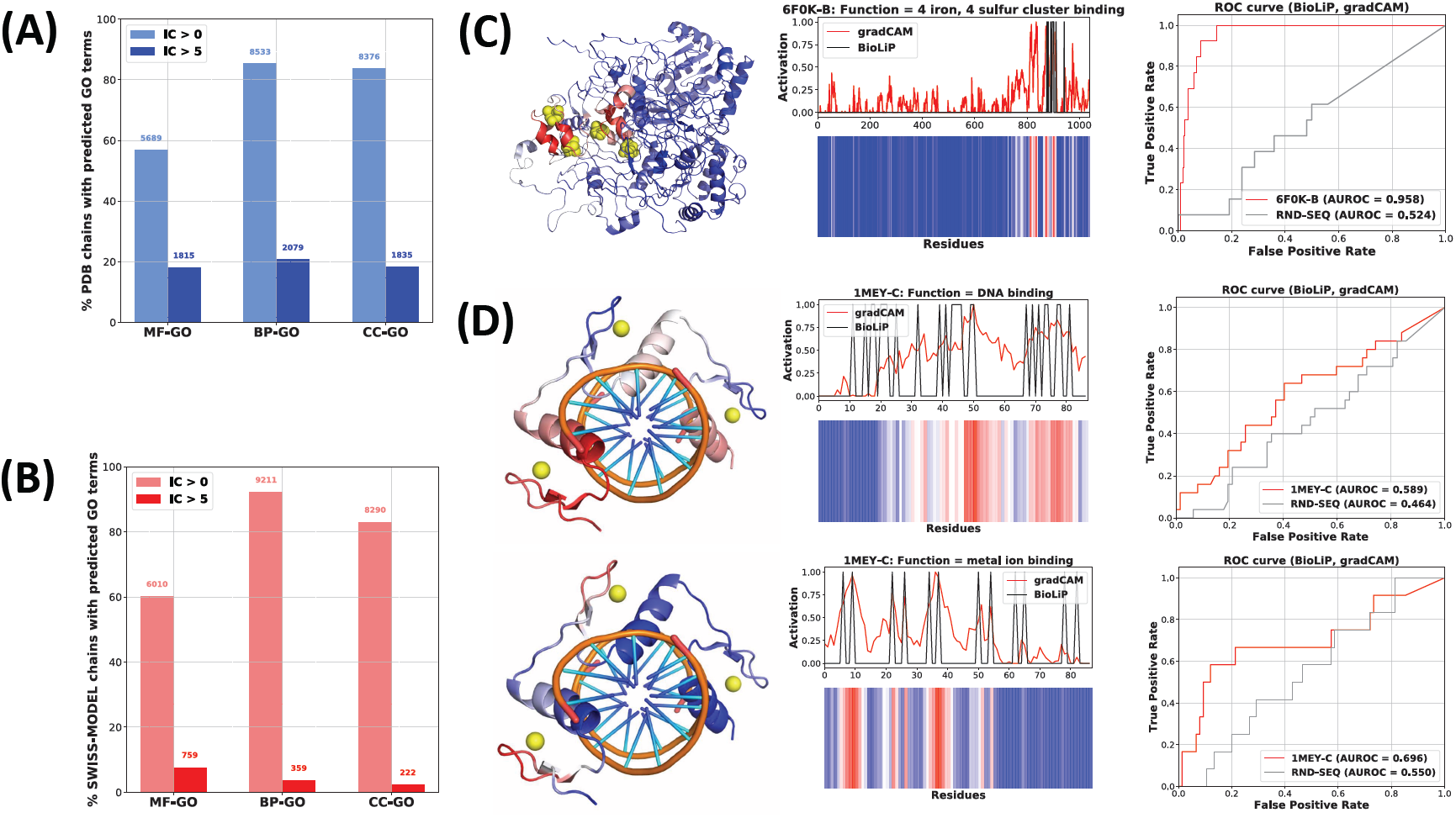
Predicting and mapping function to unannotated PDB & SWISS-MODEL structures. Percentage/number of PDB chains (A) and SWISS-MODEL chains (B) with MF-, BP- and CC-GO terms predicted by our method; the number of specific GO term predictions (with IC > 5) are shown in blue and red color for PDB and SWISS-MODEL chains, respectively. (C) An example of a Fe-S-cluster-containing hydrogenase protein (PDB Id: 6F0K), found in *Rhodothermus marinus*, with missing GO term annotations in SIFTS (“unannotated”). The PDB chain lacks annotations in databases used for training our model and predicted to bind 4Fe4S iron-sulfur cluster with high confidence score; The predicted grad-CAM profile significantly overlaps with lignad-binding sites of 4Fe4S obtained from BioLiP as shown by the ROC curve. (D) grad-CAM profiles for predicted *DNA binding* and *metal ion binding* functions mapped onto the 3D structure of an “unannotated” zinc finger protein (PDB Id: 1MEY) found in *Escherichia coli*; the corresponding ROC curves show the significant overlap between grad-CAM profile and binding sites obtained from BioLiP.

## Discussion

In this work, we described a novel deep learning-based method for predicting protein function from both protein sequences and contact map representations of protein structures. Our method, trained on protein structures from PDB and SWISS-MODEL, is very efficient, and it is capable of predicting both GO terms and EC numbers of proteins and improves over state-of-the-art sequencebased methods on the majority of function terms. Features learned from protein sequences by the LSTM-LM and from contact maps by the GCN lead to substantial improvements in protein function prediction accuracy, which could enable novel protein function discoveries. It is important to note here that although high quality sequence alignment is often sufficient to transfer fold or structure information ^59^, sequence alignments are not straightforward to use to transfer function (as evidenced by the poor performance of the CAFA-like BLAST benchmark) due to the need for different thresholds for different functions, partial alignments and domain structure, protein moonlighting, and neofunctioalization ^27, 29, 60^. Thus, one important advantage of our method is that it makes progress towards function predictions that go beyond homology-based transfer by extracting local sequence and global structure features ^27^.

By comparing function prediction performance on both DMPFold and Rosetta-predicted structures and their corresponding experimentally determined structures we demonstrate that our method has a robust high denoising power. Our method’s robustness to structure prediction error indicates that it can also be reliably used in predicting functions of proteins with computationally inferred structures. The ability to use predicted structures opens a door for characterizing many proteins lacking experimentally determined 3D structures and the contents of many databases with available predicted structures (e.g., homology-based SWISS-MODEL^13^, and ModBase^20^ can be used for expanding the train set and improving predictive power of the model). The more extensive use of homology models allowed by the denoising properties of our network architecture will be a subject of future study.

While this paper mainly focuses on introducing efficient and accurate function prediction models, it also provides a means of interpreting prediction results. We demonstrate, on multiple different GO terms, that the *DeepFRI* grad-CAM identifies structurally-meaningful protein regions encompassing functionally relevant residues (e.g., ligand-binding residues). For some PDB chains, the accuracy of the *DeepFRI* grad-CAM in identifying binding residues is quite remarkable, especially given the fact the model is not principally designed to predict functional residue location and that the ligand-binding information was not given to the model *a priori*. However, the main disadvantage of considering this to be a *site-specific function prediction method* is in the multiple different meanings of grad-CAMs. Specifically, for some GO terms related to “binding”, grad-CAMs do not necessarily identify binding residues/regions; instead, they identify regions of residues that are conserved among the sequences annotated with the same function. The most interesting example demonstrating this property is *maltose binding (GO:1901982)* (see Supplementary Fig. 7). In this example, the salient residues are far from the residues binding maltose in the 3D structure; but, by looking at a few non-redundant PDB sequences annotated with *maltose binding*, we find that the grad-CAM always identifies the same residues that are conserved across the sequences. These can be explained with the fact that any neural network, including ours, would always tend to learn the most trivial features that lead to the highest accuracy ^61, 62^.

After the culmination of much effort, two key problems in computational biology, *protein structure prediction* and *protein function prediction*, are linked together by the described methods. Deep learning coupled with an increasing amount of available sequence and structural data has the potential to meet the annotation challenges posed by ever increasing volumes of genomic sequence data, offering several new methods for interpreting protein biodiversity across our expanding molecular view of the tree of life.

## Acknowledgements

RJX is funded by NIH, JDRF and Center for Microbiome Informatics and Therapeutics. RB is funded by NSF 1728858-DMREF and NSF 1505214 - Engineered Proteins. TK is partly funded by the Polish National Agency for Academic Exchange grant PPN/PPO/2018/1/00014 to TK. RB, VG, PDR, DB, CC and JKL are supported by Simons Foundation funding to the Flatiron Institute. KC is partly supported by Samsung AI and Samsung Advanced Institute of Technology.

## Competing Interests

The authors declare that they have no competing financial interests.

## Methods

### Construction of contact maps

We collect 3D atomic coordinates of proteins from the Protein Data Bank (PDB) [1]. As the PDB contains extensive redundancy in terms of both sequence and structure, we remove identical and similar sequences from our set of annotated PDB chains. We create a non-redundant set by selecting PDB chains that are not identical to any other PDB chain in the set. To do so, we first cluster all PDB chains (for which we were able to retrieve contact maps) by *blastclust* at 100% sequence identity (i.e., number of identical residues out of the total number of residues in the sequence alignment). Then, from each cluster we select a ‘’representative” PDB chain as a PDB chain which is annotated (i.e., has at least one GO term in at least one of the 3 ontologies) and which is of high quality (has a high-resolution structure). In addition to PDB structures, we also obtained homology-based predicted structures from the SWISS-MODEL repository [2]. We include only annotated SWISS-MODEL structures (i.e., having at least one GO term in at least one of the 3 GO ontologies) in our training procedure. We remove similar SWISS-MODEL sequences in the same way as with PDB chains. Including SWISS-MODEL structures lead to 5-fold increase in the number of training samples (see **Table S1**) and also in larger coverage of more specific GO terms (see **Supplementary Fig. 4**).

To construct contact maps, we use the alpha-carbon, C_α_, atom type and consider two resides to be in contact if the distance between their corresponding C_α_ atoms is less than 10Å. We refer to this type of contact maps as “CA-CA”. We have also considered two other criteria for contact map construction. Two residues are in contact: 1) if the distance between any of their atoms is less than 6.5 Å (we refer to this type of contact maps as “*ANY-ANY”*) and 2) if the distance between their Rosetta neighbor atoms is less than sum of the neighbor radii of the amino acid pair (we refer to this type of contact maps as “*NBR-NBR”*). Rosetta neighbor atoms are defined as the beta-carbon (C_ß_) for all amino acids except glycine where the alpha-carbon is used. An amino acids neighbor-radius describes a potential interaction sphere that would be swept out by the side amino acid side chain as it samples all possible conformations. Neighbor-neighbor contact maps are therefore more indicative of side-chain-side-chain interactions than maps C_α_ - C_α_. To avoid training the model on protein chains with long sequences, we only construct contact maps for chains > 60 and < 1200 residues. We have also experimented with different cut-off thresholds for C_α_ - C_α_ and *ANY-ANY* contact maps. We found that our method produced similar results when trained on these contact maps with CA-CA (C_α_ - C_α_ with 10Å) producing slightly better results (see **Supplementary Fig. 1**).

While some recent works show better structure-prediction performance when using explicit distance matrices for predicting protein structure, here the contact maps are an input and only a means of composing graph convolutions (discrete paths for convolutions over sequence-features), and thus and there is no evidence that distances would inherently perform better than contacts in this setting. Our experiments show that additional detail in the protein structure representation (via more complex or multi-modal approaches to the contact graph) are not likely to lead to substantive performance or interpretability gains.

### Function annotations of PDB & SWISS-MODEL chains

In the training of our models we use two sets of function labels: 1) Gene Ontology (GO) [3] terms and 2) enzyme commission (EC) numbers [4]. GO terms are hierarchically organized into 3 different onotologies - molecular function (MF), biological process (BP) and cellular component (CC). We train our models to predict GO terms separately for each ontology. The summary of GO identifiers as well as EC numbers for each PDB and SWISS-MODEL chain were retrieved from SIFTS [5] (Structure integration with function, taxonomy and sequence) database and UniProt Knowledgebase databases, respectively.

SIFTS transfers annotation to PDB chain level via residue-level mapping between UniProtKB and PDB entries. All the annotation files were retrieved from SIFTS database (2019/06/18) with PDB release 24.19 and UniPortKB release 2019.06. We consider annotations that are: 1) not electronically inferred (in figure captions/legends, we refer to those as “EXP”), specifically, we consider GO terms with the following evidence codes: EXP, IDA, IPI, IMP, IGI, IEP, TAS and IC and 2) electronically inferred (in figure captions/legends, we refer to those as “IEA”).

Furthermore, we focus only on specific MF-, BP- and CC-GO terms that have enough training examples from the non-redundant training set (see the section above). That is, we select only GO terms that annotate > 30 (for MF and CC) and > 50 (for BP) non-redundant PDB/SWISS-MODEL chains. We retrieved enzyme classes for sequences and PDB structures from the lowest level (most specific level) of EC tree. The number of GO terms and EC classes in each ontology is represented in **Table S1**.

In our analyses, we differentiate GO terms based on their specificity, expressed as Shannon Information Content (IC) [6]:

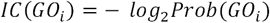

where, *Prob(GO*_*i*_) is the probability of observing GO term *i* in the UniProt-GOA database (*Prob(GO*_*i*_*)=n*_*i*_/*n*, where *n*_*i*_ – number of proteins annotated with GO term *i* and n – total number of proteins in UniProt-GOA). Infrequent GO terms (i.e., more specific) have higher *IC* values.

### Predicted structures

The initial set of benchmark structures used here was Jane and Dave Richardson’s ‘’Top 500’’ dataset [7]. It is a set of hand curated, high quality (the top 500 best), protein structures that were chosen for their fit to their completeness, how well they fit the experimental data, and lack of high energy structural outliers (bond angle and bond length deviations[8]). This set has been used in the past for fitting Rosetta energy/score terms and numerous other structural-bioinformatics validation tasks. Unfortunately, the structures in this set lacked sufficient annotations (many of these structures were the results of structural genomics efforts and had no, or only high level, annotations in GO and EC). Accordingly, we choose an additional 300 sequences from the PDB. These additional high-quality benchmark structures were chosen by taking 119K chains with function annotations and filtering them with the PISCES Protein Sequence Culling Server [9] with the following criteria:

Sequence percentage identity: <= 25

Resolution: 0.0 ∼ 2.0

R-factor: 0.2

Sequence length: 40 ∼ 500

Non- X-ray entries: Excluded

CA-only entries: Excluded

Cull PDB by chain

That left us with 1606 SIFTS annotated chains from which we randomly selected 300. These proteins were then excluded from all phases of model training. The performance of our method on this set of PDB chains is shown in **Fig. 2A** in the main manuscript. In **Supplementary Fig. 3** we demonstrate capabilities of our method in denoising these predicted structures.

### Convolutional neural network

Convolutional neural networks (CNNs) have shown tremendous success in extracting information from sequence data and making highly accurate predictive models. Their success can be attributed to convolutional layers with highly reduced number of learnable parameters which allow multi-level and hierarchical feature extraction. In the last few years, a large body of work has been published covering various applications of CNNs, such as prediction of protein functions [10] and subcellular localization [11], prediction of effects of noncoding-variants [12] and protein fold recognition [12]. Here we use the CNN-based *DeepGOPlus* method [10] in our comparison study. We describe this architecture in more detail in the Supplementary Material.

We represent a protein sequence with *L* amino acid residues as a feature matrix ***X*** = [***x***_1_ … ***x***_*L*_] ∈ {0,1}^*L*×*c*^, where c = 26 dimensions (25 residues plus the gap symbol) are used as a one-hot indicator, ***x***_*i*_ ∈ {0,1}^*c*^, of the amino acid residue at position *i* in the sequence. This representation is fed into a convolution layer which applies a one-dimensional convolution operation with a specified number of kernels (weight matrices or filters), *f*_n_, of certain length, *f*_l_, and all outputs are then transformed by the rectified linear activation function (*ReLU*), which sets values below 0 to 0, i.e., *ReLU*(x) = max(x, 0). This is followed by a global max pooling layer and a fully connected layer with *sigmoid* activation function for predicting probabilities of GO terms or EC enzyme classes.

In the first convolution layer, we used 16 CNN layers with *f*_n_ = 512 filters of different lengths (see Supplementary Material). After concatenating the outputs of the CNN layers, we obtain *L*×8192 dimensional feature map for each sequence. Using filters of variable lengths ensures extraction of complementary information from protein sequences. The second layer has |GO|^1^ number of units for GO terms (or |EC| for EC) classification.

### LSTM language model for learning residue-level features

We use an approach similar to *Bepler & Berger* [13]. We train a LSTM language model on ∼10,000,000 sequences sampled from the entire set of sequences from *Pfam*. The sequences are represented using 1-hot encoding (see above). The LM architecture is comprised of two stacked forward LSTM layers with 512 units each (see **Fig. 1** in the main manuscript). The LSTM LM model is trained for 5 epochs using ADAM optimizer with learning rate *lr* = 0.001 and batch size of 128.

The residue-level features, extracted from the final LSTM layer’s hidden states, ***H***^*LM*^, are combined together with 1-hot representation of sequences, ***X***, through learnable non-linear mapping:

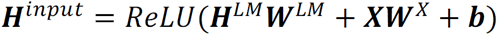

where ***H***^*input*^ is the final residue-level feature representation passed to the fist GCN layer, ***H***^(0)^ = ***H***^*input*^ (see the equation below). We refer to this stage of our method as a feature extraction stage. The parameters, ***W***^*LM*^, ***W***^*X*^ and ***b*** are trained together with the parameters of the GCN. All the parameters of the LSTM LM are frozen during the training. We choose this strategy because it more efficient (i.e., instead of fine tuning the huge number of the LSTM-LM parameters together with GCN parameters, we only tune, ***W***^*LM*^, ***W***^*X*^and ***b*** parameters while keeping the parameters of the LSTM-LM fixed).

### Graph Convolutional Network

Graph Convolutional Networks (GCNs) have proven to be powerful methods for extracting features from data that are naturally represented as one or more graphs [14]. Here we experiment with the notion that GCN are a suitable method for extracting features from proteins by taking into account their graph-based structure of amino acids represented by contact maps. We propose our model based on the work of *Kipf & Welling* [15]. A protein graph can be represented by an adjacency matrix (also termed contact map), ***A*** ∈ ℝ^***L***×***L***^, encoding connections between its *L* residues, and a residue-level feature matrix, ***X*** ∈ ℝ^*L*×*c*^.

We explore different residue-level feature representations including the one-hot encoding representation of residues as in the CNN (*c=26*), LSTM language model (*c=512*, i.e., the output of the LSTM layers), and no sequence features (to be able to run GCN, in this case, feature matrix is substituted with an diagonal identity matrix, i.e., ***X*** = ***I***_***L***_.

The graph convolution takes both adjacency matrix, *A*, and residue-level embeddings from the previous layer, 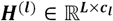 and outputs the residue-level embeddings in the next layer, **H**^(*l*+1)^ ∊ 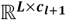:

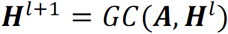

where ***H***^(0)^ = ***H***^*input*^, and *c*_*l*_ and *c*_*l*+*1*_ are residue embedding dimensions for layers *l* and *l*+*1*, respectively. Concretely, we use the formulation of *Kipf & Welling* [15]:

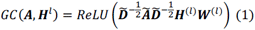

where ***Ã*** = ***A*** + ***I***_*L*_ is the adjacency matrix with added self-connections represented by the identity matrix 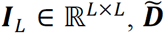 is the diagonal degree matrix with entries 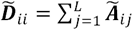 and 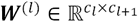 is a trainable weight matrix for layer *l*+*1*.

To normalize residue features after each convolutional layer the adjacency matrix is first symmetrically normalized, hence the term 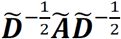 Equation (1) updates features of each residue by a weighted sum of features of the residue in its one-hop neighborhood (adding self-connections ensures that the residue’s own features are also included in the sum).

Given that we are classifying individual protein graphs with different number of residues, we use several layers, *N*_*l*_*=3*, of graph convolutions. The final protein representation is obtained by first concatenating features from all layers into a single feature matrix, i.e., 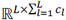 and then by performing a global pooling layer after which we obtain a fixed vector representation of a protein structure, 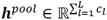. The global pooling is obtained by a sum operator over ***L*** residues:

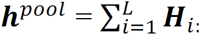

We then use a fully connected layer with *ReLU* activation function for computing hidden representation from the pooled representation. This is then followed by a fully connected layer which used for mapping the hidden representation from the previous layer to a |GO| x 2 output; that is, two activations for ach GO term. These activations are transformed by a *softmax* activation outputting the positive and negative probability for each GO term/EC number; (i.e., the final layer outputs probability vector 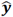 of dimension |GO| x 2 (|EC| x 2 for EC numbers) for predicting positive and negative probabilities of GO terms (EC numbers).

### Model training and hyperparameters

To account for imbalanced label problem, both CNN and GCN are trained to minimize *weighted binary cross-entropy* cost function that gives higher weights to GO term with fewer training examples:

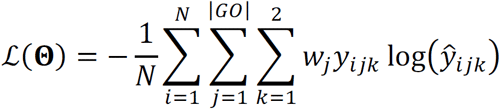

where Θ is the set of all parameters in all layers to be learned; 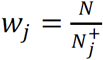 is class weight for function *j*, with 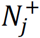 being the number of positive examples associated with function *j*; *N* is the total number of samples and |GO| is the total number of functions (i.e., GO terms); *y*_*ijk*_ is the true binary indicator for sample *i* and function *j* (i.e., *y*_*ij*1_ = 1, *y*_*ij*2_ = 0, if sample *i* is annotated with function *j*) and *ŷ*_*ij*1_ is the predicted probability that sample *i* is annotated with function *j*. In the inference phase, we say we predict GO terms/ EC numbers if the positive probability is > 0.5.

All hyperparameters are determined through grid search based on the model’s performance on the validation set. The validation set is comprised of ∼10% randomly chosen samples from the training set. To avoid overfitting, we use early stopping with *patience=5* (i.e., we stop training if validation loss does not improve in 5 epochs). We use ADAM optimizer [16] with learning rate *lr* = 0.0001, *β*_1_= 0.95 and *β*.= 0.95 and batch size of *64*. The default number of epochs is 200. Both GCN and CNN are implemented to deal with variable length sequences, by performing sequence/contact map padding. The entire method is implemented using *Keras* deep learning library.

### Training and test set construction

We partition the non-redundant set composed of both PDB and SWISS-MODEL sequences into train, validation and test sets, with ratio 80%/10%/10% such that for each function we have at least 20 training examples with direct experimental evidence (GO evidence code EXP) validated annotations (15 SWISS-MODEL and 5 PDB) and at least 2 test examples (1 SWISS-MODEL and 1 PDB). The test set is chosen to be no more than 40% sequence identical to the training set. We use cd-hit tool [17] to split the sequences into training, validation and test sets. We perform experiments with different thresholds. In all our experiments we trained the model using both EXP and IEA GO annotations (see **Supplementary Fig. 2**), but the test set, composed of only experimentally annotated PDB & SWISS-MODEL chains (EXP), is always kept fixed. See **Table S1**.

We examine the performance of our method when trained only on PDB, SWISS-MODEL and both PDB & SWISS-MODEL contact maps; we also investigate training on only EXP and both EXP and IEA function labels (see **Supplementary Fig. 2**). In all our experiments the final results are averaged over 100 bootstraps of the test set.

### Temporal holdout validation

We also evaluate the performance of our method by using temporal holdout validation approach similar to CAFA [18]. The temporal holdout approach ensures a more “realistic” scenario where function predictions [19] are evaluated based on recent experimental annotations. We used GO annotations retrieved from SIFTS [5] from two time points, version 2019/06/18 (we refer to this as SIFTS-2019) and version 2020/01/04 (we refer to this as SIFTS-2020), to construct our temporal holdout test set. We form the test set from the PDB chains that did not have any annotations in SIFTS-2019 but gained annotations in SIFTS-2020. To increase the GO term coverage, we focus on the PDB chains with both “EXP” and “IEA” evidence codes. We obtain 4,072 PDB chains (out of which 3,115 have sequences < 1200 residues). We use our model (trained on SIFTS-2019 GO annotations) to predict functions of these newly annotated PDB chains. We evaluated our predictions against the annotations from SIFTS-2020.

The results for MF-, BP- and CC-GO terms are show in **Supplementary Fig. S17**. We also show a few examples of the PDB chains with correctly predicted MF-GO terms by our method, for which both *BLAST* and *DeepGO* are not able to make any significant predictions.

### Residue-level annotations

We use a method based on Gradient-weighted Class Activation Map (grad-CAM) [20] to localize function predictions on a protein structure (i.e., to find residues with highest contribution to a specific function). Grad-CAM is a class-discriminative localization technique that provides visual explanations for predictions made by CNN-based models. Motivated by its success in image analysis, we use grad-CAM to identify important, function-specific residues in a protein structure.

In a grad-CAM approach, we first compute the contribution of each filter, *k*, in the last convolutional layer to the prediction of function label *l* by taking derivative of the output of the model for function *l, y*^l^, with respect to feature map ***F***_*k*_ ∈ ℝ^*L*^ over the whole sequence of length *L*:

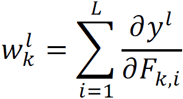

where 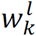 represent the importance of feature map *k* for predicting function *l*, obtained by summing the contributions from each individual residue. Finally, we obtain the function-specific heat-map in a residue space by making the weighted sum over all feature maps in the last convolutional layer:

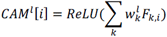

where *ReLU* function ensures that only features with positive influence on the functional label are preserved; *CAM*^*l*^[*i*] - indicates the relative importance of residue *i* to function *l*. The advantage of grad-CAM is that it does not require re-training or changes in the architecture of the model which makes is computationally efficient and directly applicable to our models. See **Supplementary Figs. 8-16** representing grad-CAM mapped onto 3D structure of PDB chains with known ligand-binding information.

### Data availability

Our training, validation and test data splits are available from: https://users.flatironinstitute.org/~vgligorijevic/DeepFRI_data/

### Code availability

Source code for training the *DeepFRI* model, together with neural network weights are available for research and non-commercial use at: https://github.com/flatironinstitute/DeepFRI Web service of our method is available at: https://beta.deepfri.flatironinstitute.org/

## Footnotes

1 |GO|, |EC| - denotes the number of GO term, EC numbers in the set.

## Notes

### Competing Interest Statement

The authors have declared no competing interest.

### Summary of Updates

New results on Swiss-Model structures added. New figures with class activation map added. Figures 2, 3, and 4 revised. Author list updated. Supplementary material updated.

