## Supplementary Material for "Structure-Based Protein Function Prediction using Graph Convolutional Networks"

**This PDF file includes:**

Model architectures
Figs. S1 to S17
Tables S1
References

**Other Supplementary Materials for this manuscript include the following:**

Excel Tables S1 to S2:
S1_DeepFRI_unannotPDB_predictions.xls
S2_DeepFRI_unannotSWISS_predictions.xls

**CNN architecture** (using *Keras*[1] syntax)

**Input:** 1-hot encoding of sequence: (n_samples, L, 26).

- `1D CNN layer x 16 (# filters = {512, 512, 512, 512, 512, 512, 512, 512, 512, 512, 512, 512, 512, 512, 512, 512}, length = {8, 16, 25, 32, 40, 48, 56, 64, 72, 80, 88, 96, 104, 112, 120, 128}, l2_reg = 2e-4)`
- `Concatenate layer`
- `BatchNormalization`
- `Activation (ReLU)`
- `Dropout (0.3)`
- `GlobalMaxPooling`
- `Dropout (0.6)`
- `Dense (dim = |GO| or dim = |EC|)`
- `Activation (sigmoid)`

Optimization: loss = `binary_crossentropy`; optimizer = `Adam` (lr = 0.0005, β1 = 0.95, β2 = 0.99); `batch_size` = 64; `epochs` = 100; `EarlyStopping` (patience=5).

[1] https://keras.io/

**LSTM-LM architecture**

**Input:** 1-hot encoding of sequence: (n_samples, L, 26).

- `LSTM (dim = 512, return_sequences=True, kernel_constrain=MinMax(-2.0, 2.0), recurrent_constrain=MinMax(-2.0, 2.0))`
- `LSTM (dim = 512, return_sequences=True, kernel_constrain=MinMax(-2.0, 2.0), recurrent_constrain=MinMax(-2.0, 2.0))`
- `TimeDistributed(Dense(26))`
- `Activation (softmax)`

Optimization: loss = `categorical_crossentropy`; optimizer = `Adam` (lr = 0.001, β1 = 0.99, β2 = 0.99); `batch_size` = 128; `epochs` = 5.

**GCN architecture**

**Input:** 1-hot encoding of sequence, S = (n_samples, L, 26); Normalized contact maps, A = (n_samples, L, L); Pre-trained LSTM-LM.

- `Sequence embedding:`
  - `SeqEmbedding_1 = Dense (dim = 128, use_bias=False)(S)`
  - `SeqEmbedding_2 = Dense (dim = 128, use_bias=False)(LSTM-LM(S))`
  - `Add()(SeqEmbedding_1, SeqEmbedding_2)`
  - `Activation (ReLU)`
- `GCN layer (dim = 256, use_bias=False, l2_reg = 2e-4)`
- `Activation (ReLU)`
- `GCN layer (dim = 256, use_bias=False, l2_reg = 2e-4)`
- `Activation (ReLU)`
- `GCN layer (dim = 512, use_bias=False, l2_reg = 2e-4)`
- `Activation (ReLU)`
- `Concatenate layer (all GCN layers)`
- `GlobalSumPooling`
- `Dropout(0.3)`
- `Dense (dim = 1024)`
- `Activation (ReLU)`
- `|GO| X Dense (dim = 2)`
- `Activation (softmax)`

Optimization: loss = `categorical_crossentropy`; optimizer = `Adam` (lr = 0.001, $\beta 1$ = 0.99, $\beta 2$ = 0.99); `batch_size` = 64; `epochs` = 200; `EarlyStopping` (patience=5).

**Evaluation metrics**

We evaluate the performance of our method using both *protein-level* and *residue-level* metrics as follows:

1) *Protein-level evaluation:* we measure the performance of our method in a) predicting functions for a particular protein (*protein-centric*) and b) predicting protein associated with a particular GO/EC term (*term-centric*). To this end, we use two measures first proposed in CAFA[18]:

   a) *Protein*-centric F-max obtained by finding the maximum of $F_1$ score over thresholds $t \in [0,1]$:

$$F_{max} = \max_t \left\{ \frac{2 \cdot AvgPr(t) \cdot AvgRc(t)}{AvgPr(t) + AvgRc(t)} \right\}$$

   where, precision is averaged over all proteins, *m(t)*, for which we predict at least one term: $AvgPr(t) = \frac{1}{m(t)} \sum_{i=1}^{m(t)} pr_i(t)$, whereas recall is averaged over all proteins, *n*: $AvgRc(t) = \frac{1}{n} \sum_{i=1}^{n} rc_i(t)$. For a given target protein, $i$, and some value of threshold $t \in [0,1]$, the precision and recall are computed as:

$$pr_i(t) = \frac{\sum_f I(f \in P_i(t) \wedge f \in T_i)}{\sum_f I(f \in P_i(t))}$$
$$rc_i(t) = \frac{\sum_f I(f \in P_i(t) \wedge f \in T_i)}{\sum_f I(f \in T_i)}$$

   where *f* is a GO/EC term, $T_i$ is a set of known GO/EC terms for protein *i* (for MF-GO, BP-GO and CC-GO we propagated annotations up to the root term), and $P_i(t)$ is a set of predicted GO/EC terms with score ≥ t, $I(\cdot)$ – the indicator function.

   b) *Term-centric* area under the Precision-Recall curve (AUPR), where precision and recall for each term *f* are computed as:

$$pr_f(t) = \frac{\sum_i I(f \in P_i(t) \wedge f \in T_i)}{\sum_i I(f \in P_i(t))}$$
$$rc_f(t) = \frac{\sum_i I(f \in P_i(t) \wedge f \in T_i)}{\sum_i I(f \in T_i)}$$

   For each term, *f*, we compute PR curve using the sliding window method (i.e., across all threshold values of $t \in [0,1]$) and then we compute AUPR using the trapezoid rule. Even though in same cases we report AUPR values for each individual GO term (e.g., **Fig. S6**), in most cases we report the AUPR performance under *micro*- and *macro*- averaging; *micro*-averaged PR curve is computed by first vectorizing the protein–function predicted scores and known binary annotations (i.e., flattening protein-function matrices), and then computing the PR curve using the sliding window method (e.g., **Fig. 2A**, **B**). The area under the PR curve, obtained by applying trapezoid rule, is known as *micro*-AUPR. *macro*-AUPR (*macro*-AUPR) is computed by first computing the AUPR for each function separately, and then averaging these values across all functions.

2) *Residue-level* evaluation: for each individual protein and its predicted MF-GO term, we measure the ability of our method in predicting binding/active sites. This measure can only be computed for the minority of proteins with detailed site specific annotations; here we rely on this site specific annotation encoded in the BioLiP database [21]. For a given protein of $L$ residues, we construct a ligand-binding binary profile (retrieved from BioLiP), $\boldsymbol{s} \in \{0, 1\}^L$, indicating residues known to bind a specific ligand (e.g. ATP); i.e., $\boldsymbol{s}_i = 1$ if residue $i$ is a ligand-binding residue, $\boldsymbol{s}_i = 0$ otherwise. For the same protein and its corresponding predicted function (e.g., *ATP binding* (GO:0005524)), we compute a real-valued grad-CAM profile from our pre-trained *DeepFRI* method, $\hat{\boldsymbol{s}} \in [0,1]^L$, indicating the functional importance of each residue. To show how well the grad-CAM profile recovers known binding sites, we compute the area under the ROC curve, representing the values of sensitivity (recall) for a given 1-specificity (false positive rate), using the sliding threshold approach; we then compute area under the ROC curve (AUROC) using the trapezoid rule [22]. See **Figs. S8-S16** for examples of ROC curves for different GO terms.

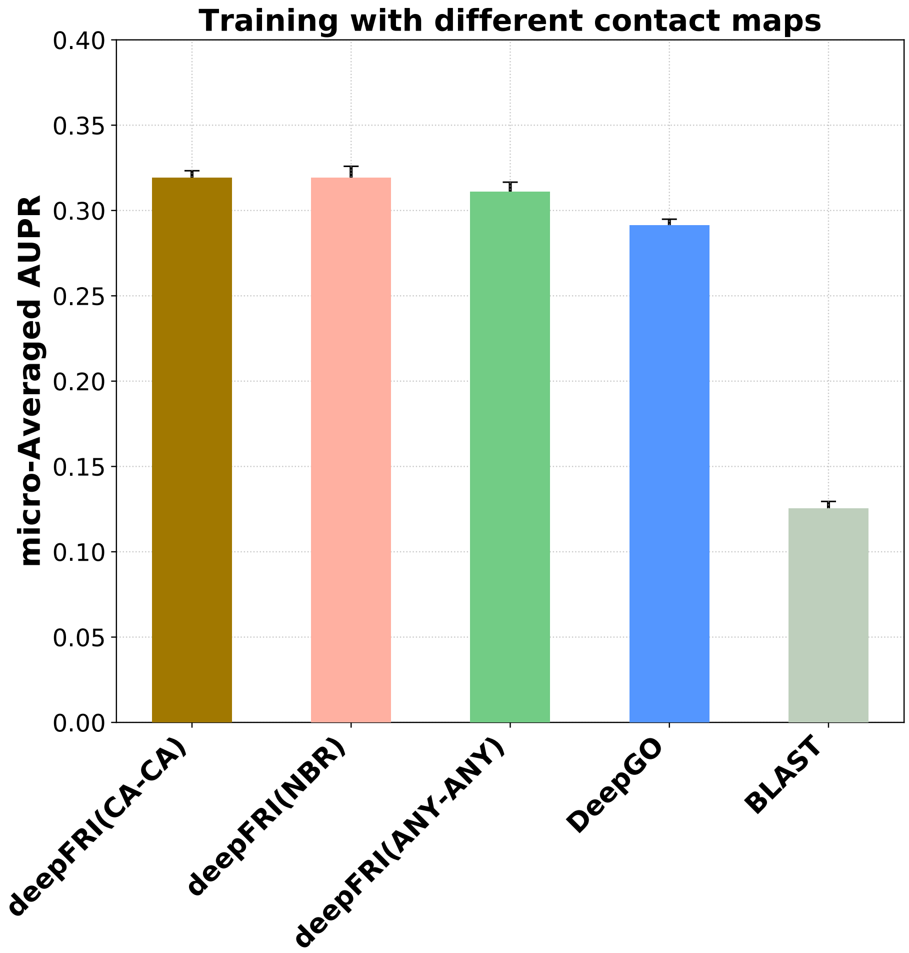


**Fig. S1.** Performance of our method in comparison to state-of-the-art CNN (*DeepGO*) and *BLAST* baseline trained on different contact maps. The model is trained on proteins with experimental (EXP) MF-GO annotations. The results are averaged over 100 bootstraps of the test set.

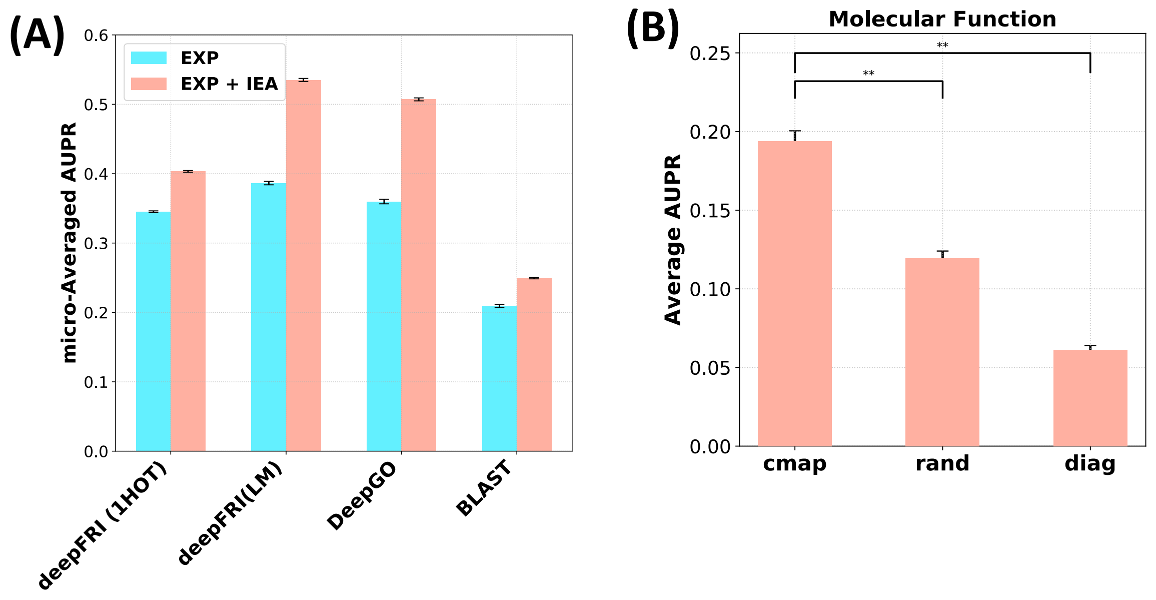


**Fig. S2**. (A) Performance of our method in comparison to state- of-the-art CNN (*DeepGO*) and *BLAST* baseline trained with different sequence features ("1HOT" - 26-dimensional binary one-hot encoding of residues, "LM" - features from the pretrained LSTM Language Model) and trained using protein chains with experimental (EXP) only and electronically inferred (EXP+IEA) MF-GO annotations. Test proteins used to compose this metric were annotated with EXP evidence codes. (B) Performance of our model trained on CA-CA contact maps with experimental (EXP) annotations and evaluated on test set composed on CA-CA contact maps ("cmap"), generated contact maps with random contacts ("rand") and contact maps with not contacts except for self-loops ("diag"). Asterisks indicate where the performance of our method trained on CA-CA contact maps is significantly better than its' performance trained on "rand" and "diag" (rank-sum p-value < 0.001).

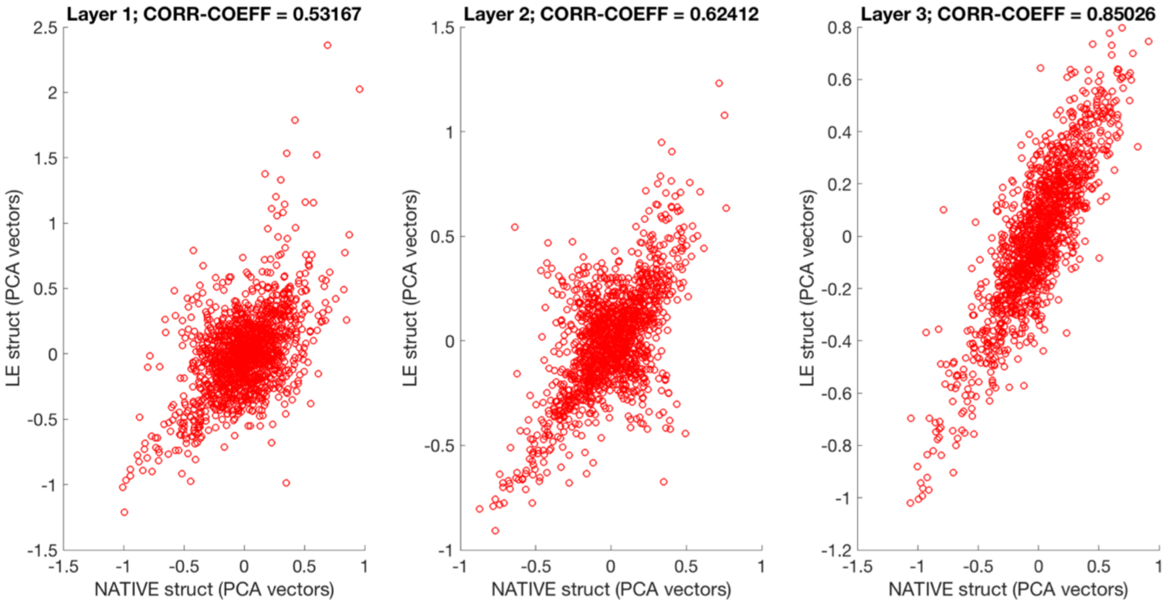


**Fig. S3**. De-noising Rosetta contact maps. Here we plot features from using NATIVE contact maps as input vs. the features derived from using Rosetta-predicted contact maps. We show these features for the 1st, 2nd and 3rd layer of graph convolution, and note that the features extracted from the third layer of the GCN exhibit strong NATIVE-Predicted correlations, providing strong evidence that our model is tolerant to even significant error in structure predictions, and can effectively denoise structure predictions.

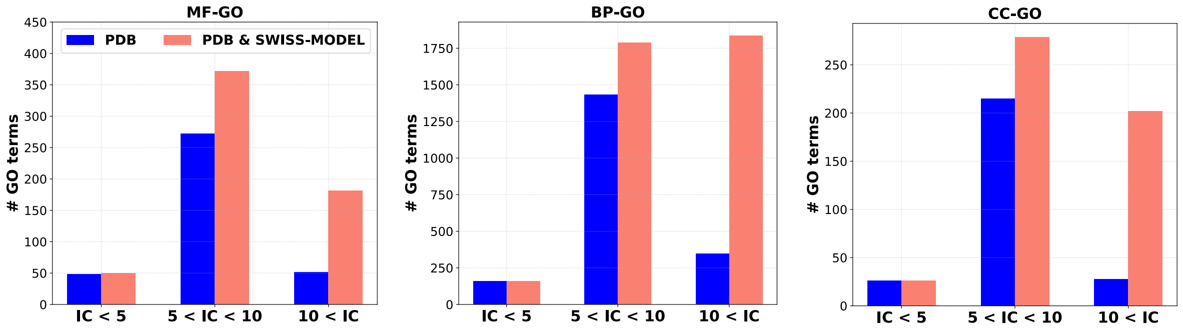


**Fig. S4**. We stratify MF-, BP- and CC-GO term into different groups based on their specificity, expressed as Information Content (IC), see methods, and show the number of each term in each category encompassed by both the PDB-only-trained (blue) and the PDB-and-SWISS-MODEL trained models (red).

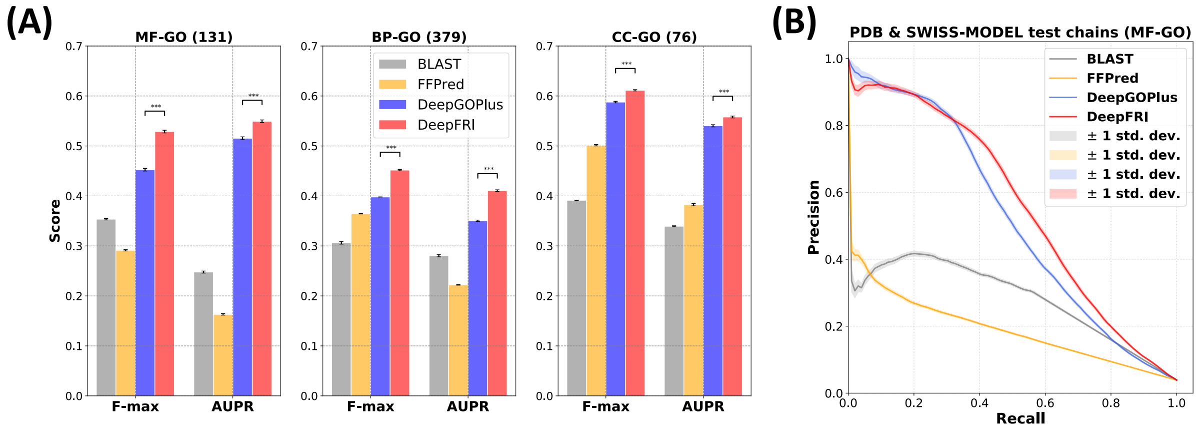


**Fig. S5**. (A) F-max and AUPR scores, summarized over all proteins and GO terms, respectively, computed on the test set comprised of PDB and SWISS-MODEL chains chosen to have < 40 % sequence identity to the sequences in the training set. The numbers in brackets indicate the number of GO terms in different ontologies that are common to all 4 methods; Asterisks indicate where the performance of *DeepFRI* is significantly better than *DeepGOPlus* (rank-sum pval < 0.001). The total number of GO terms for *DeepFRI* and *DeepGOPlus* is much higher and it is shown in Fig 2C in the main manuscript. (B) Micro-average precision-recall curves for each method for MF-GO terms. The curves are averaged over 100 bootstraps of the test set.

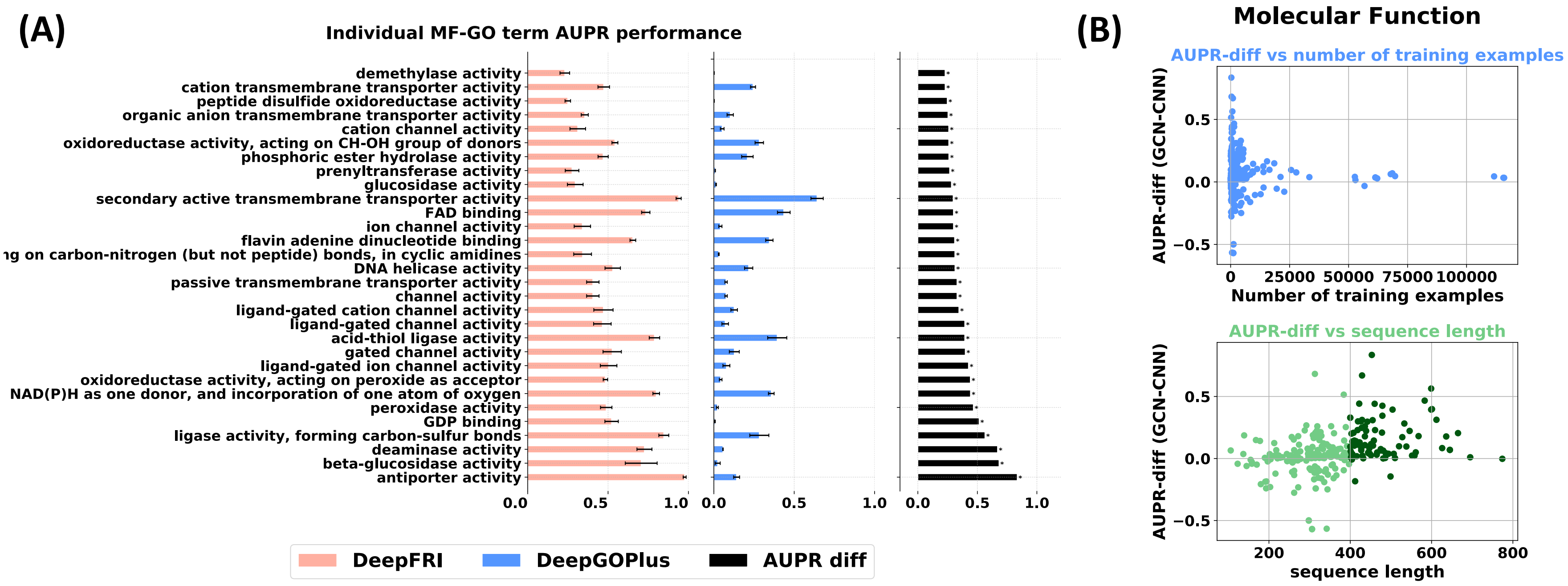


**Fig. S6.** Experiments on experimentally annotated PDB chains show difference in the performance of our method vs. CNN in predicting individual MF-GO terms. For each MF-GO term we show *DeepFRI* and *DeepGOPlus* performance measured by the AUPR averaged over 100 bootstraps of the test proteins. The third panel shows the difference in the performance of these two methods. Only top 30 MF-GO terms on which *DeepFRI* > *DeepGOPlus* are shown.

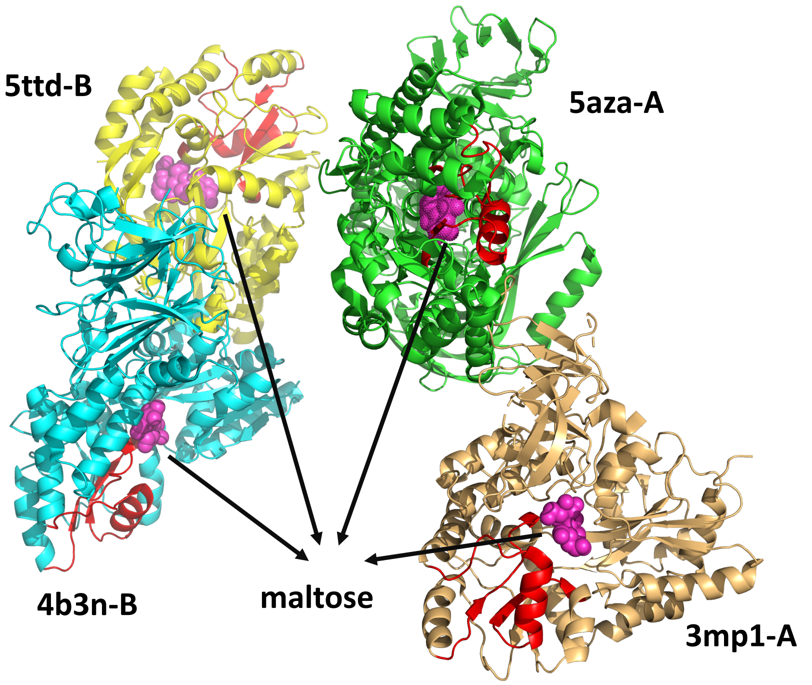


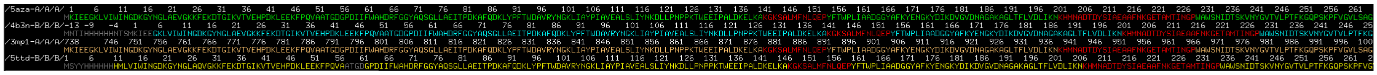

**Fig. S7**. Class Activation Map of "maltose binding" PDB chains. The salient regions, shown in red, are away from maltose-binding regions. However, they are consistent (evolutionary conserved) across different PDB chains.

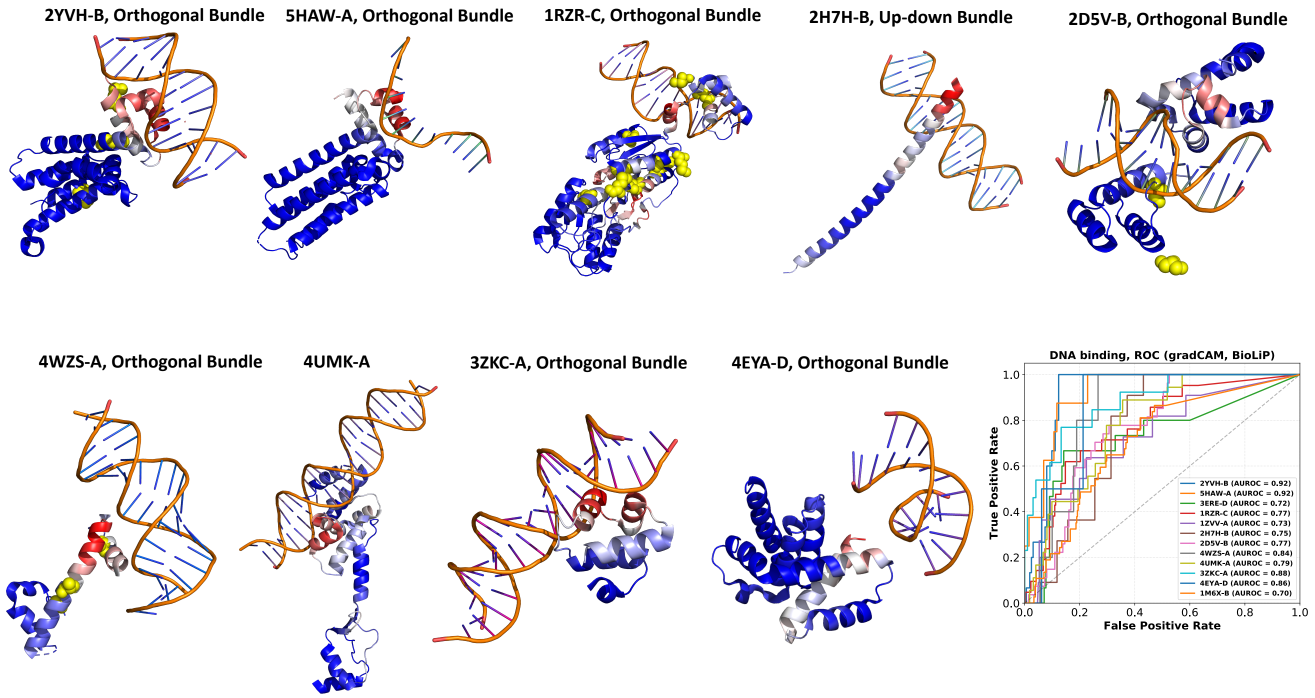


**Fig. S8. Grad-CAM for "DNA binding" (GO:0003677) mapped onto the 3D structure of the test PDB chains annotated with "DNA binding".** Their corresponding ROC curves measuring the overlap between the grad-CAM profile and DNA binding sites (retrieved from BioLiP database) are shown in the bottom right corner.

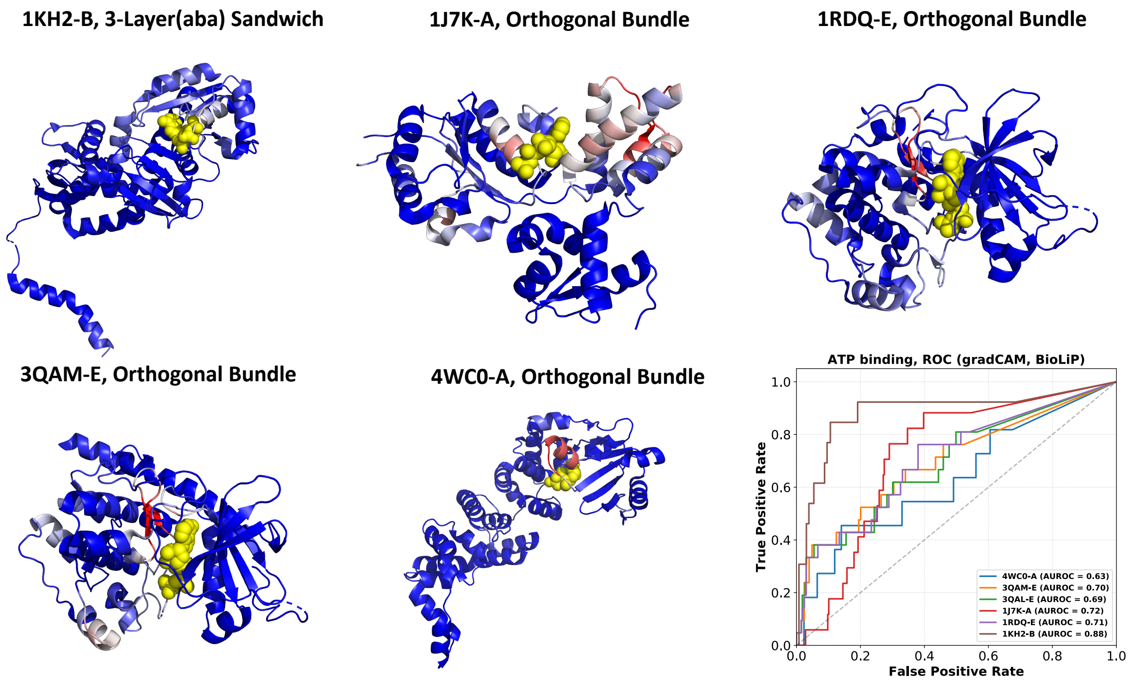


**Fig. S9. Grad-CAM for "ATP binding" (GO:0005524) mapped onto the 3D structure of the test PDB chains annotated with "ATP binding".** Their corresponding ROC curves measuring the overlap between the grad-CAM profile and ATP binding sites (retrieved from BioLiP database) are shown in the bottom right corner.

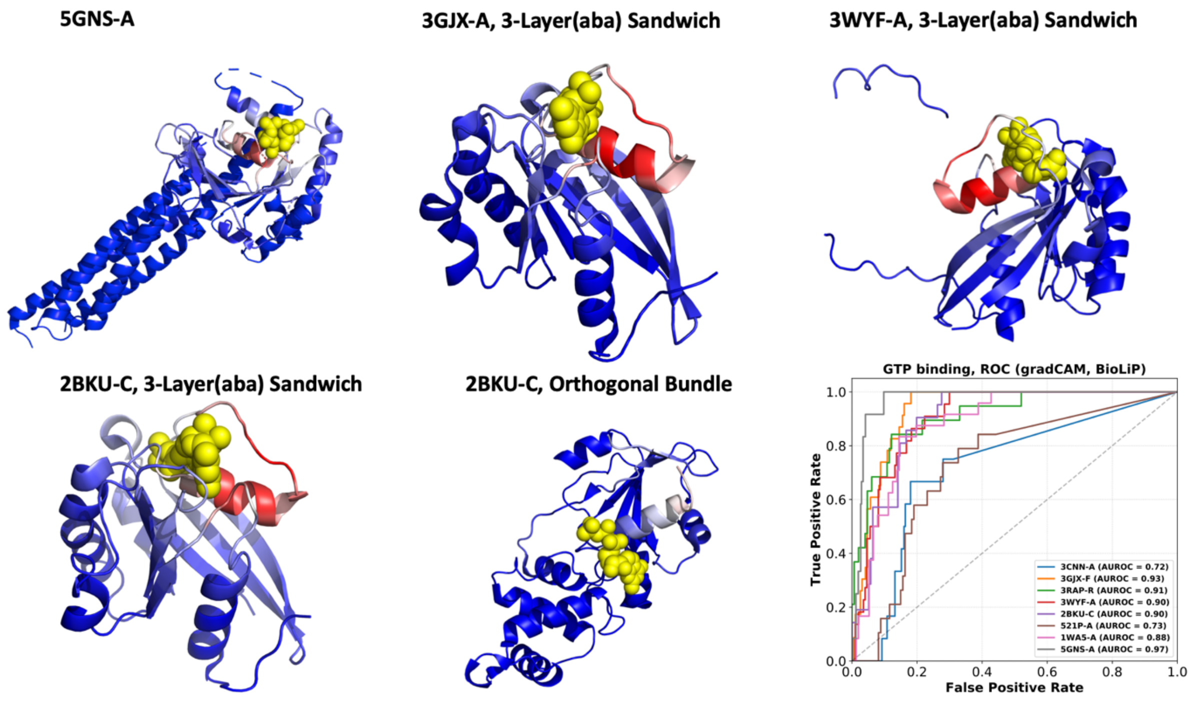


**Fig. S10. Grad-CAM for "GTP binding" (GO:0005525) mapped onto the 3D structure of the test PDB chains annotated with "GTP binding".** Their corresponding ROC curves measuring the overlap between the grad-CAM profile and GTP binding sites (retrieved from BioLiP database) are shown in the bottom right corner.

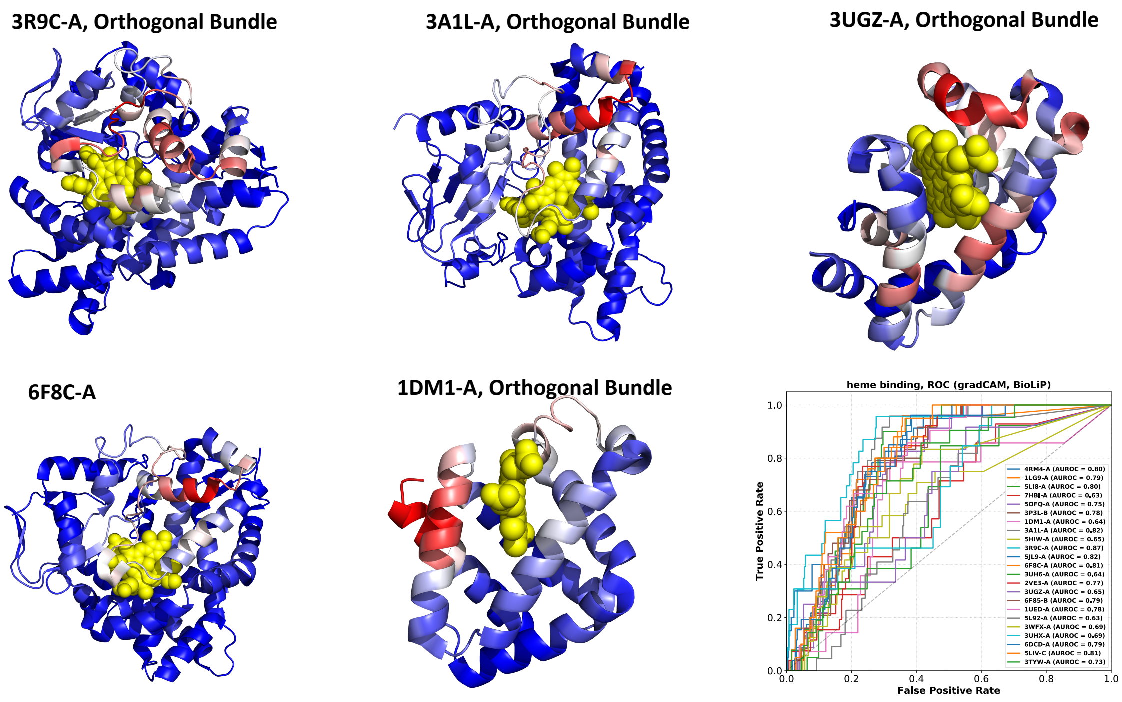


**Fig. S11. Grad-CAM for “heme binding” (GO:0020037) mapped onto the 3D structure of the test PDB chains annotated with “heme binding”.** Their corresponding ROC curves measuring the overlap between the grad-CAM profile and HEM binding sites (retrieved from BioLiP database) are shown in the bottom right corner.

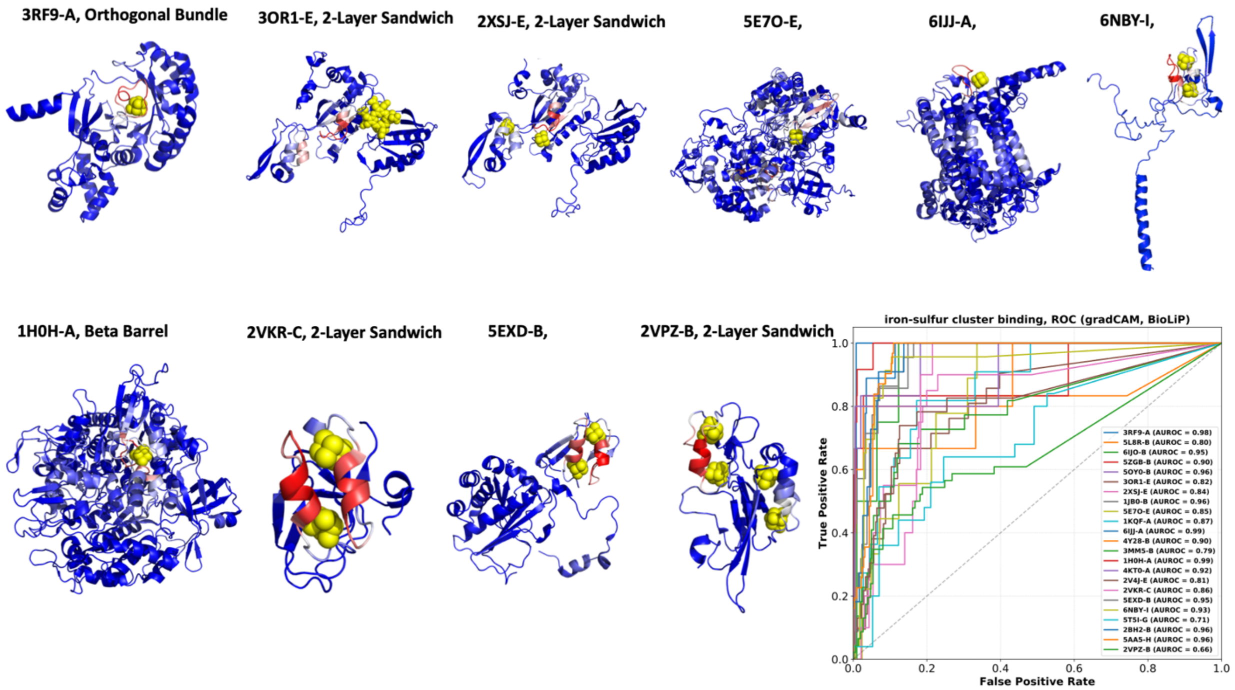


**Fig. S12. Grad-CAM for "iron-sulfur cluster binding" (GO:0051536) mapped onto the 3D structure of the test PDB chains annotated with "iron-sulfur cluster binding".** Their corresponding ROC curves measuring the overlap between the grad-CAM profile and SF4 binding sites (retrieved from BioLiP database) are shown in the bottom right corner.

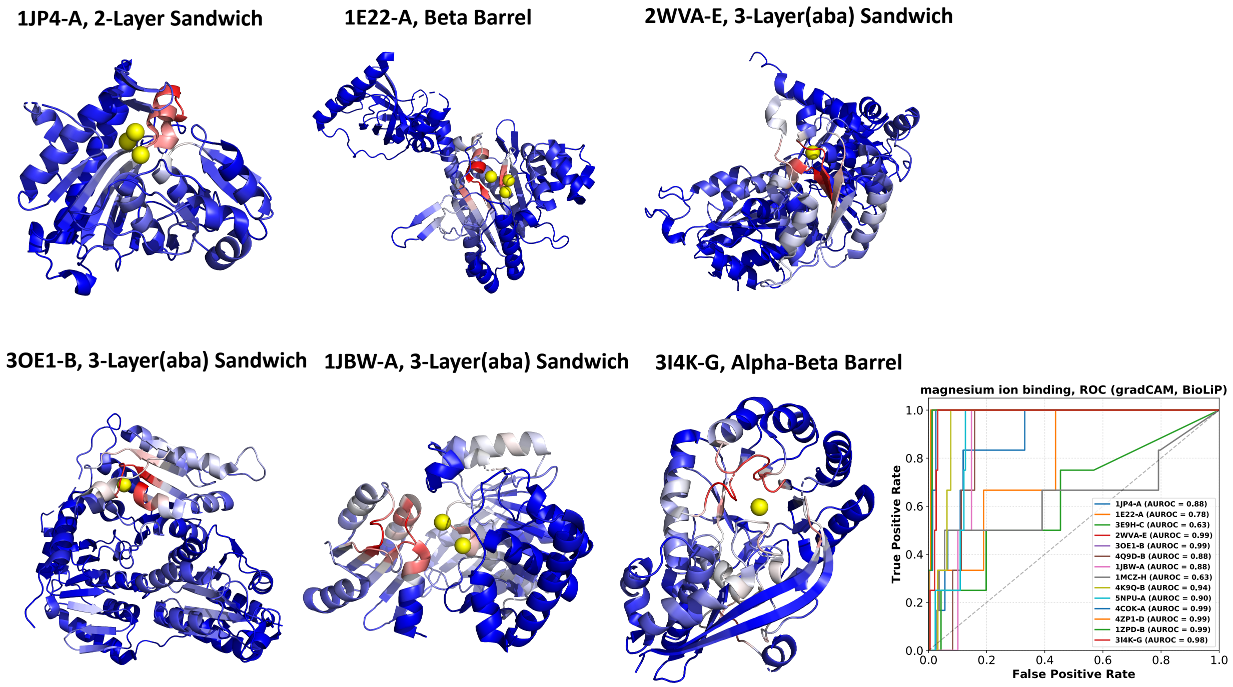


**Fig. S13. Grad-CAM for “magnesium ion binding” (GO:0000287) mapped onto the 3D structure of the test PDB chains annotated with “magnesium ion binding”.** Their corresponding ROC curves measuring the overlap between the grad-CAM profile and MG binding sites (retrieved from BioLiP database) are shown in the bottom right corner.

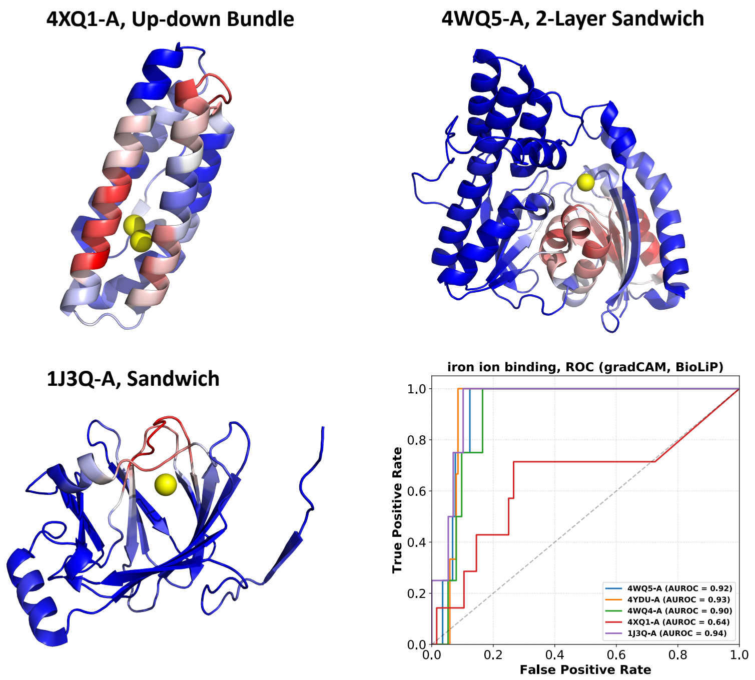


**Fig. S14. Grad-CAM for "iron ion binding" (GO:0005506) mapped onto the 3D structure of the test PDB chains annotated with "iron ion binding".** Their corresponding ROC curves measuring the overlap between the grad-CAM profile and FE binding sites (retrieved from BioLiP database) are shown in the bottom right corner.

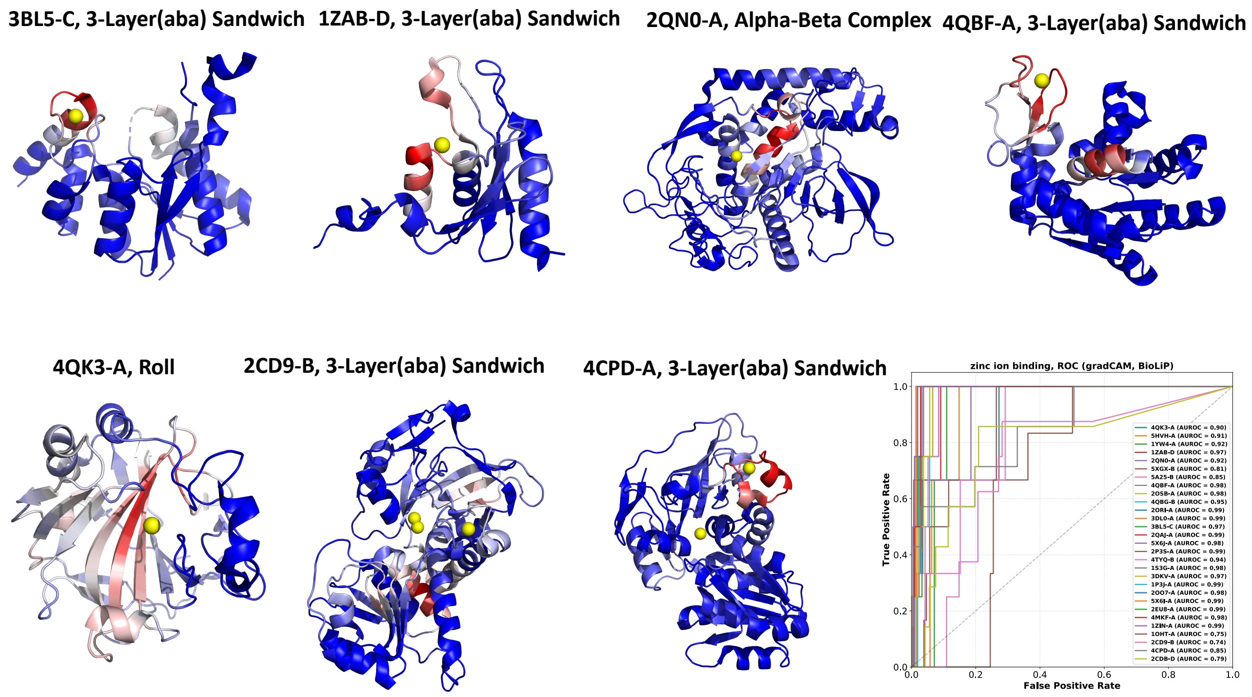


**Fig. S15. Grad-CAM for "zinc ion binding" (GO:0008270) mapped onto the 3D structure of the test PDB chains annotated with "zinc ion binding".** Their corresponding ROC curves measuring the overlap between the grad-CAM profile and ZN binding sites (retrieved from BioLiP database) are shown in the bottom right corner.

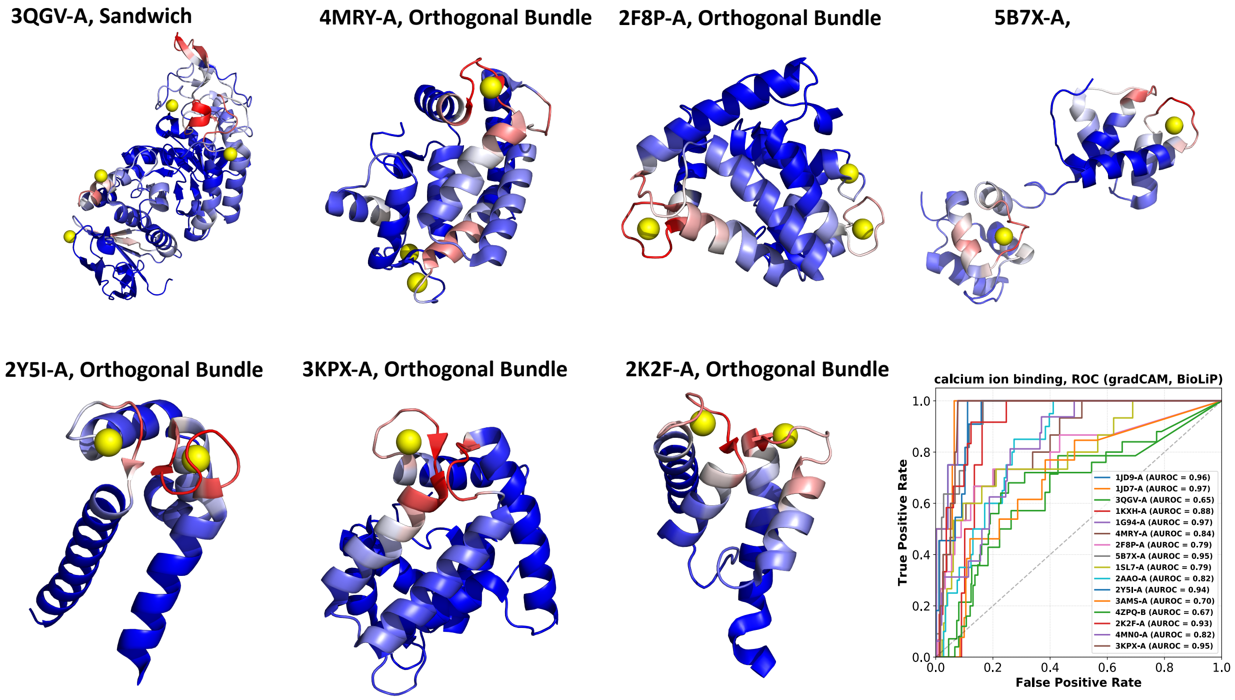


**Fig. S16. Grad-CAM for "calcium ion binding" (GO:0005509) mapped onto the 3D structure of the test PDB chains annotated with "calcium ion binding".** Their corresponding ROC curves measuring the overlap between the grad-CAM profile and CA binding sites (retrieved from BioLiP database) are shown in the bottom right corner.

(A)

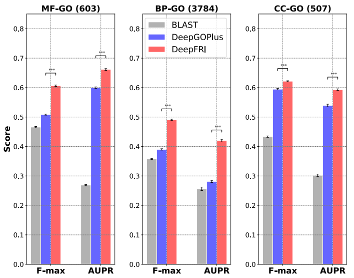


(B)

| PDB-chain | GO term | GO name | BLAST | DeepGO | DeepFRI |
|---|---|---|---|---|---|
| **6AGZ-B** | GO:0010181 | FMN binding | 0.35 | 0.01 | 1.00 |
| **6Q72-E** | GO:0070279 | vitamin B6 binding | 0.00 | 0.09 | 1.00 |
| **6E6B-A** | GO:0005509 | calcium ion binding | 0.45 | 0.98 | 0.99 |
| **6OR8-A** | GO:0051539 | 4 iron, 4 sulfur cluster binding | 0.00 | 0.35 | 0.98 |
| **6NHG-E** | GO:0051537 | 2 iron, 2 sulfur cluster binding | 0.00 | 0.25 | 0.98 |
| **6RTD-A** | GO:0020037 | heme binding | 0.00 | 0.21 | 0.87 |
| **6GTJ-A** | GO:0005507 | copper ion binding | 0.26 | 0.10 | 0.87 |
| **6ONS-A** | GO:0051539 | 4 iron, 4 sulfur cluster binding | 0.47 | 0.10 | 0.86 |
| **6IWP-A** | GO:0008289 | lipid binding | 0.41 | 0.06 | 0.85 |
| **6JO6-A** | GO:0051536 | iron-sulfur cluster binding | 0.00 | 0.01 | 0.79 |
| **6OHK-A** | GO:0010181 | FMN binding | 0.37 | 0.02 | 0.79 |
| **6AIR-A** | GO:0051536 | iron-sulfur cluster binding | 0.00 | 0.27 | 0.78 |
| **6S0O-D** | GO:0005507 | copper ion binding | 0.34 | 0.04 | 0.73 |
| **6DE9-A** | GO:0051536 | iron-sulfur cluster binding | 0.00 | 0.02 | 0.69 |
| **6NCO-A** | GO:0008289 | lipid binding | 0.00 | 0.06 | 0.56 |
| **6S6X-J** | GO:0051539 | 4 iron, 4 sulfur cluster binding | 0.00 | 0.06 | 0.52 |

(C)

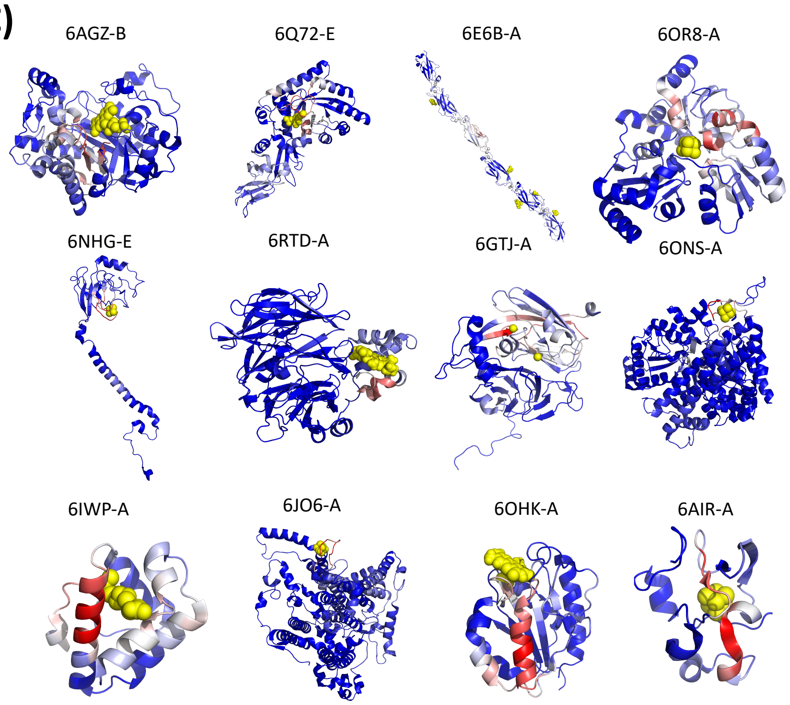


**Fig. S17. Temporal holdout validation.** (A) Average protein-centric F-max and term-centric AUPR values of our method in comparison to the CAFA-like *BLAST* baseline and *DeepGOPlus* applied on temporal holdout test PDB chains. The values are averaged over 100 bootstraps of the test set; asterisks indicate where the performance of *DeepFRI* is significantly better than the performance of *DeepGOPlus* (rank-sum p-value < 0.01); (B) PDB chains correctly annotated with our method (prediction score > 0.5) with very low *BLAST* and *DeepGOPlus* prediction scores; the low scores indicate the inability of *BLAST* and *DeepGOPlust* to correctly infer their GO terms. These PDB chains were selected because they have ligand-binding information in *BioLiP* that allows us to validate our Class Activation Mapping identification of functional sites on protein sequences and structures; (C) Grad-CAM profile mapped onto 3D stricture of the proteins in the Table.

| Data | Annotations | Train | Test | Validation | # terms |
|---|---|---|---|---|---|
| PDB | MF | 31,254 | 2,855 | 7,926 | 603 |
| | BP | 29,065 | 1,993 | 7,343 | 3,784 |
| | CC | 17,396 | 1,535 | 4,366 | 507 |
| | EC | 10,994 | 1,725 | 1,888 | 545 |
| SWISS-MODEL | MF | 149,321 | 4,148 | 37,209 | 603 |
| | BP | 152,101 | 4,448 | 37,893 | 3,784 |
| | CC | 118,118 | 2,947 | 29,395 | 507 |
| | EC | 56,425 | 11,132 | 9,858 | 507 |

**Table S1**. Table showing the number of PDB & SWISS-MODEL chains in train, test and validation sets in each GO branch and EC classification system. Test chains are chosen to have ≤ 40% sequence identity to the chains in the training set.
